# Regulation of *N*-glycosylation efficiency by eukaryotic oligosaccharyltransferase

**DOI:** 10.1101/2025.09.06.674603

**Authors:** Marium Khaleque, Beatrice Tropea, Ojas Singh, Danila Elango, Shulei Liu, Joel D. Allen, Maddy L. Newby, Abigail S. L. Sudol, Jacob Willcox, John Butler, Patrick Duriez, Francesco Forconi, Ivo Tews, Amanda S. Nouwens, Max Crispin, Elisa Fadda, Benjamin L Schulz

**Affiliations:** School of Chemistry and Molecular Biosciences, The University of Queensland, St Lucia, QLD 4072, Australia; Hamilton Institute, Maynooth University, Maynooth, Ireland; Department of Chemistry, Maynooth University, Maynooth, Ireland; School of Biological Sciences, University of Southampton, Southampton, United Kingdom; Centre for Cancer Immunology, University Hospital Southampton, Southampton, United Kingdom; School of Cancer Sciences, Faculty of Medicine, University of Southampton, Southampton, United Kingdom

## Abstract

Oligosaccharyltransferase (OST) catalyses the key step of *N*-glycosylation, transferring immature *N*-glycans to select Asn residues in nascent proteins in the endoplasmic reticulum (ER). Asn are more likely to be glycosylated in the context of a “glycosylation sequon”, N-x-S/T (x≠P), but not every sequon is glycosylated. Tight positive and negative regulation of site-specific *N*-glycosylation across the glycoproteome is essential because some Asn require glycosylation for productive protein folding or function, while others must remain unglycosylated. Despite its importance, the underlying bases for this regulation are illdefined. Here we characterise the molecular determinants regulating OST catalytic efficiency. We identify preferred substrates of OST, and show that the local sequence determines glycosylation efficiency by fine-tuning binding to an extended groove in OST along an eight residue stretch centered at the Asn acceptor. Tight control of OST activity is achieved through a ‘switch’, set ‘ON’ when a peptide binds with high complementarity to the groove, triggering release of the assisting base Glu350 for catalysis. Based on analysis of buried sequons, we identify sequence characteristics associated with inherently non-preferred acceptor substrates for OST, and show these have a high propensity for local secondary structure incompatible with the optimal conformation of a peptide bound to OST. Finally, we validate these mechanisms by examining the determinants of efficient glycosylation at a variably glycosylated Asn in a set of human Fabs. These mechanistic and functional insights show how local protein sequence controls glycosylation occupancy with immediate implications in secretory protein function and evolution, pathogen immunity and epistasis, and in engineering protein sequences for the desired degree of site-specific glycosylation.

## Introduction

*N-*glycosylation is an essential biological process present in all domains of life that results in the covalent functionalization of selected Asn sidechains with a glycan structure^1–3^. The initial stage of *N-*glycosylation occurs co- or post-translationally on a nascent protein entering the secretory pathway, where select Asn are covalently modified with glycan transferred from a lipid-linked oligosaccharide (LLO) donor by the oligosaccharyltransferase (OST). In eukaryotes the efficiency of glycosylation of Asn is dramatically enhanced if they are located within a **N**-x-S/T sequon, where x cannot be Pro^4,5^, while the sequon for the bacterial OST is longer, D/E-x’-**N**-x-S/T, where neither x’ or x can be Pro and where an acidic residue is required at the −2 position^6^. The structural bases for the glycosylation sequon are specific binding sites in OST: a pocket that accommodates and hydrogen bonds with the OH of the Ser/Thr at the +2 position of the acceptor peptide; and in the bacterial OST an Arg that forms a salt bridge with the Asp/Glu at the −2 position. The “sequon” provides sufficient binding affinity to allow modification of the Asn, but in a simple motif that can be readily gained and lost through evolution in diverse sequence and structural contexts in substrate proteins.

Occupancy of Asn in *N-*glycosylation sequons is dictated by substrate recognition and by the catalytic activity of OST, which is not equally efficient at all sites, resulting in different degrees of *N*-glycosylation of sequons, or macroheterogeneity^7^. More specifically, about 2/3rds of Asn in sequons appear to be efficiently glycosylated^8–10^, while the rest remain essentially unmodified, and rarely Asn in ‘non-canonical’ sequons with Cys or Val at the +2 aa instead of Ser/Thr can be glycosylated^11^. In most cases, glycosylation at a specific Asn is not critical to protein structure and function, but in others tight control of OST catalytic activity is required. Indeed, OST activity is likely optimised such that glycosylation is efficient at Asn where *N-*glycosylation is required for protein folding efficiency, quaternary association, structural stability, and/or biological function. Conversely, OST should not efficiently glycosylate Asn where it would interfere with folding or with key biomolecular interactions, such as in the protein interior, in close proximity to binding sites or at protein-protein interfaces. These functional constraints to OST substrate recognition and activity exist in the context of a large and diverse glycoproteome, with 100,000s of *N*-glycosylation sites in 10,000s of glycoproteins in a typical eukaryote, with a corresponding vast variety of potential acceptor sequences and contexts. Some features of efficiently modified sequons have been previously identified based on the analysis of limited data sets or within the context of specific substrates, such as sequons containing Thr rather than Ser at the +2 position^9,12^, and with small hydrophobic amino acids at the +1 position^8,13–15^. However, these traits are not strict determinants of *N*-glycosylation occupancy, and do not explain how OST is able to modify a broad variety of sequences while maintaining such complex substrate selectivity.

Here we combined molecular and cell biology, mass spectrometry glycoproteomics, molecular dynamics (MD) simulations, free energy calculations, and data analytics to characterise the molecular determinants that regulate the substrate specificity and catalytic efficiency of eukaryotic OST. We first monitored changes in site-specific *N*-glycosylation occupancy in yeast cell wall proteins as a function of different glycosylation stresses. We found that *N*-glycosylation sequons were differentially impacted by the various glycosylation stresses, which allowed us to identify the subset of sites preferred as OST substrates. Molecular dynamics (MD) simulations showed that OST substrate specificity is controlled by interactions ‘beyond the sequon’ along a stretch of residues between +4 and −4 around the target Asn. The acceptor peptide’s residues within this stretch can engage in interactions with an extended peptide binding groove in OST. These not only stabilise the binding of the substrate at the OST catalytic site, but also activate the enzyme by inducing a structural switch from its inactive resting state to a catalytically productive ON state. Free energy calculations show that when preferred substrates are bound, the ON state of the switch is energetically favoured; both ON and OFF states are energetically accessible to substrates whose glycosylation efficiency depends on glycosylation stress conditions; and only the OFF state is energetically favoured for sites that are poorly glycosylated. We describe the mechanism by which the OST switch is turned ON and OFF through interactions with the backbone of tightly bound acceptor peptides, operating independently of the amino acid sequence. Finally, we identify the sequence and structural features of poorly glycosylated Asn across the secreted proteome that explain how these sites are inefficient substrates of OST, and validate these features in a model glycoprotein to provide a full-circle understanding of the molecular determinants regulating *N*-glycosylation in eukaryotes.

## Results and Discussion

### Quantitative yeast glycoproteomics identifies the preferred acceptor substrates of OST

To understand the sequence determinants regulating OST efficiency we performed quantitative glycoproteomics to measure *N*-glycosylation site occupancy in yeast cell wall glycoproteins under two different glycan stress conditions (**Fig. 1a**). We chose these stress conditions to induce either global under-glycosylation, or site-specific under-glycosylation such that the preferred substrates of OST would remain efficiently glycosylated. To obtain global under-glycosylation, we generated glycan concentration stress by partially inhibiting the first committing step of the *N*-glycan biosynthetic pathway, the Alg7 glycosyltransferase enzyme that catalyses the addition of the first *N*-acetylglucosamine (as GlcNAc-1-P) to dolichol phosphate on the cytoplasmic side of the endoplasmic reticulum (ER) membrane. Partial inhibition of Alg7 limits flux into LLO biosynthesis, but as the remainder of the pathway is intact results in production of full length LLO substrate (Glc_3_Man_9_GlcNAc_2_) at low concentration. We predicted that in this low concentration of full length LLO, OST would efficiently transfer glycan to any normally glycosylated acceptor peptide substrate bound to the enzyme, but at globally low efficiency. To obtain site-specific under-glycosylation at non-preferred substrates, we generated glycan structural stress by inhibiting the Alg6 glucosyltransferase enzyme that catalyses addition of the first glucose to the LLO intermediate in the ER lumen. Cells lacking Alg6 accumulate a truncated glycan (Man_9_GlcNAc_2_), but at a normal concentration^16^. We predicted that with this normal concentration of truncated LLO substrate, OST would only efficiently transfer glycan to its preferred acceptor peptide substrates, while weaker acceptor peptide substrates would be inefficiently modified. We created two independent experimental systems to generate each type of glycan stress. We inhibited Alg7 with the antibiotic tunicamycin and Alg6 with a genetic knockout, or inhibited expression of the gene encoding either enzyme with a tet::off system. We titrated the concentration of tunicamycin and doxycycline to obtain equivalent overall under-glycosylation phenotypes in each type of glycan stress (for details see Supplementary Material and **Fig. S1**).

**Figure 1.**
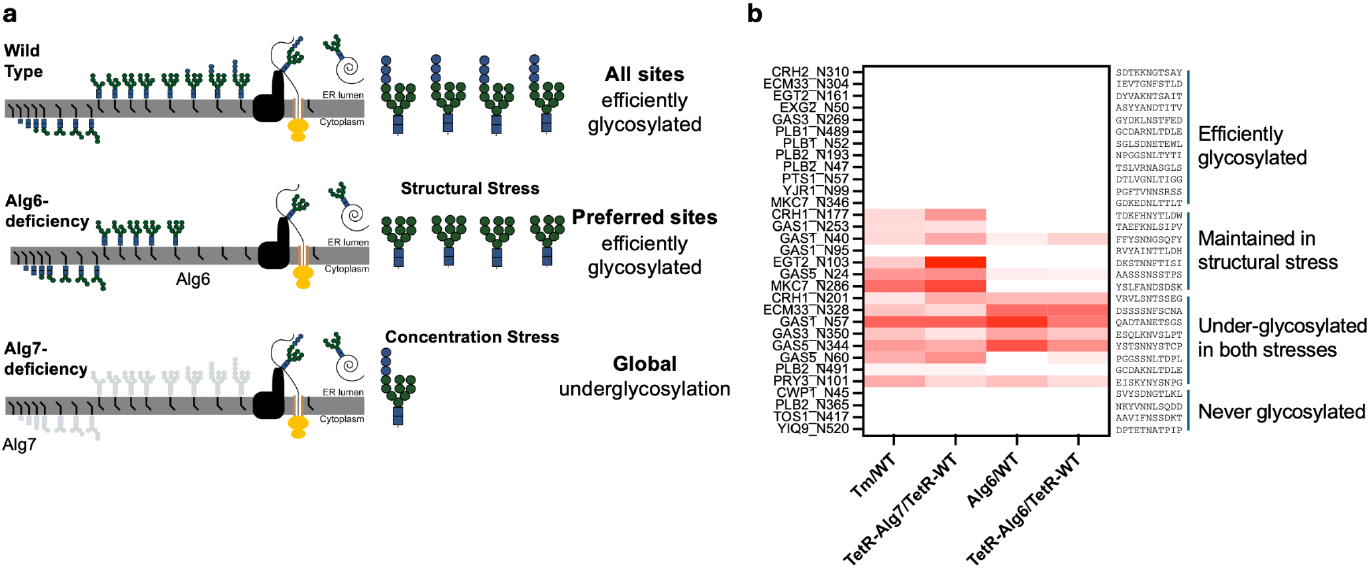
*N*-glycosylation site occupancy in glycan structural and concentration stress identifies the preferred substrates of OST. **(a)** Two experimental models of glycan structural stress (Alg6 deficiency) and glycan concentration stress (Alg7 deficiency) were established. **(b)** Log-fold change of glycosylation site occupancy in two models of glycan structural and glycan concentration stress compared to the unstressed condition; red, robustly significantly underglycosylated compared to unstressed cells (p <0.05). For details see Supplementary Material.

We grew wild type unstressed yeast cells and yeast subjected to glycan structural or concentration stress to mid-log phase, prepared cell wall protein samples, and measured site-specific glycosylation occupancy with LC-MS/MS^17,18^ (**Fig. 1b** and **Fig. S2**). As expected, in wild type cells most Asn in sequons were essentially completely modified, while the remaining few Asn in sequons were not glycosylated. The two experimental systems showed broadly consistent site-specific glycosylation occupancy in each stress condition relative to the wild-type unstressed cells. Most Asn that were fully glycosylated in wild-type cells remained fully glycosylated under both stress conditions; these sites are efficiently glycosylated by OST but without a difference in phenotype depending on the glycan stress, and had a small but non-significant enrichment in N-x-T relative to N-x-S sequons. A subset of sites were inefficiently glycosylated in both glycan stress conditions. Finally, and most interestingly, a subset of sites were inefficiently glycosylated under glycan concentration stress but remained efficiently glycosylated under glycan structural stress; consistent with our model of OST site recognition, these represent the preferred acceptor peptide substrates for *N*-glycosylation. However, no simple sequence characteristics defined these preferred acceptor substrates.

### OST acceptor substrate specificity is regulated along an extended binding groove

To characterise at an atomistic level the mechanisms regulating OST substrate specificity we selected a set of six peptide substrates representing preferred and non-preferred acceptors based on our quantitative yeast glycoproteomics data (**Fig. 1**) and characterised their binding to OST by all-atoms MD simulations within a deterministic sampling framework. The 3D models of the six complexes were based on cryo-EM data^19^ (PDB 8AGC), and included all the OST domains in direct contact with the peptide substrate, namely Stt3 and part of Ost3 (aa 211 to 345) (**Fig. 2a**).The OST models were embedded in a lipid bilayer representing the ER membrane and complemented with a lipid-linked oligosaccharide (LLO). (For details see Supplementary Material).

**Figure 2.**
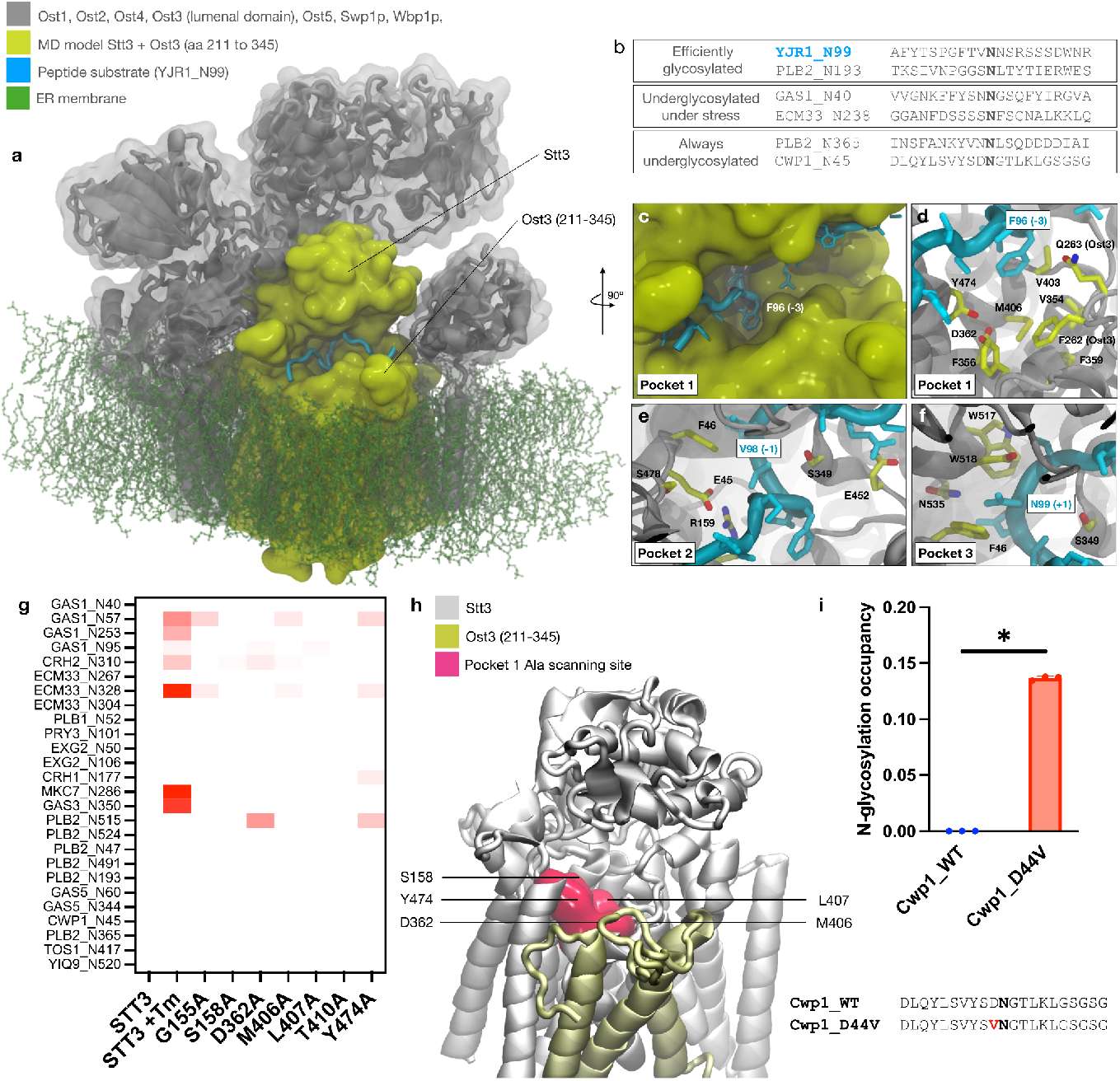
An extended acceptor peptide binding groove in OST. **(a)** 3D structure of the yeast OST built from cryo-EM structure (PDB 8AGC) with the Ost3 lumenal domain rebuilt with AlphaFold 2. The domains included in the MD model of the OST are shown in yellow within a representative (equilibrated) structure from the MD simulation in complex with the peptide YJR1_N99, shown in cyan as cartoons. The lipid bilayer representing the ER membrane in the simulation is rendered with green sticks. **(b)** Sequences of selected peptides used as substrates for the MD simulations. **(c)** Close-up view of the extended OST binding groove showing how the aromatic sidechain of Phe96 at position −3 of the acceptor peptide substrate, shown in cyan, is accommodated in a cavity (Pocket 1). **(d)** The residues of Stt3 and Ost3 lining the hydrophobic interior and the hydrophilic rim of Pocket 1 are shown as sticks with C atoms in yellow, N atoms in blue and O atoms in red. The Stt3 and Ost3 backbone is shown in grey with cartoon rendering. **(e)** Residues of Stt3, rendered with sticks, within the region identified as Pocket 2 accommodating residues at position −1 from the peptide sequon. Glu45 and Arg159 are part of the ‘switch’. **(f)** Residues of Stt3, shown with sticks, within the region identified as Pocket 3 accommodating residues at position +1 from the peptide sequon. Residue numbering follows PDB 8AGC. **(g)** Log-fold change of glycosylation site occupancy in cells expressing variant Stt3 with Ala scanning mutations across residues comprising Pocket 1 and 2, compared to wild type Stt3. Tm, tunicamycin; red, significantly underglycosylated compared to wild type Stt3 (p <0.05). For full details see Supplementary Material. **(h)** Structure of the Stt3/Ost3 complex rendered with cartoons (Stt3 white; Ost3 gold) with the residues in Pocket 1 targeted by Ala scanning shown as surface in magenta. Residues not labelled are not visible. Rendering with Visual Molecular Dynamics^21^ (VMD) (https://www.ks.uiuc.edu/Research/vmd/).**(i**) Glycosylation occupancy at Asn45 of wild type (WT) and Asp44Val variant (D44V) Cwp1. *, P<0.05. Aligned sequences of Cwp1_WT and Cwp1_D44V.

In all 3D models all peptides were rebuilt inside the OST acceptor peptide binding site with the acceptor peptide backbone from positions +3 to −3 around the sequon matching the coordinates of the substrate in the structure of the complex with bacterial OST PglB^20^ (PDB 6GXC), which in turn matches the position of the substrate in the yeast OST (PDB 8AGC), but is N-terminally extended. The peptide was then elongated residue-by-residue to cover the remaining positions up to +10/-10 aa from the target Asn, optimising the conformation of the peptide tails to prevent the formation of intrapeptide contacts. All MD simulations were run for a minimum of 1 μs of data collection for each complex, with the Cα atom of the Asn target restrained in place with a force constant of 5 kcal/molÅ^2^ to avoid unphysical displacements. For details on the method and the RMSD and RMSF analyses obtained from the MD simulations of the complexes see Supplementary Material.

The MD simulations showed that acceptor peptide substrates engaged in stable interactions with the OST far beyond the catalytic site, and that these interactions varied greatly in terms of their nature and stability, depending on the sequence of the acceptor substrate. We identified three regions within this OST binding groove that played fundamental roles in regulating binding, that we named Pocket 1, 2, and 3 (**Fig. 2c-f)**. The OST residues in these pockets can anchor the flexible peptide substrate, enhancing the stability of a catalytically productive conformation. Each of these regions is described below.

Pocket 1 accommodates the sidechain of residues at positions −3 and/or −2 of the Asn acceptor. Shaped as a cavity, lined by hydrophobic residues, with hydrophilic residues around the rim, Pocket 1 has the ideal structure and electrostatic profile to accommodate large aromatic sidechains inside the cavity, such as Phe96 at −3 in the efficiently glycosylated YJR1_N99 peptide (**Fig. 2c** and **d**), or small hydrophilic residues that can bind the rim. In contrast, the PLB2_N365 and CWP1_N45 sites are never glycosylated, and both have Tyr at −2, which MD simulations show is not an ideal residue for Pocket 1. The phenolic sidechain of Tyr cannot be easily accommodated in the hydrophobic cavity, but it is also too large to establish stable hydrophilic interactions with the rim. This lack of complementarity with Pocket 1 ultimately leads to the N-terminal section of these peptides detaching from the pocket, allowing formation of an intra-peptide hairpin structure. In the PLB2_N365 peptide this hairpin is stabilised by interaction of the Tyr at −3 with the Asp at +4 (**Fig. S5**), while in the CWP1_N45 peptide the hairpin involves the Asp at −1 and the Lys at +4. To verify the role of this Asp44-Lys49 salt bridge in stabilising a hairpin in the CWP1_N45 peptide and thereby preventing productive presentation of Asn45 to the OST active site, we expressed and purified native Cwp1 and variant Cwp1_D44V, with a mutation designed to effectively remove the salt bridge. Purified native Cwp1 was not glycosylated at Asn45 (**Fig. 2i** and **Fig. S3**), consistent with analysis of native cell wall bound Cwp1 (**Fig. 1b**). However, glycosylation at Asn45 in the Cwp1_D44V variant protein was significantly increased, with an occupancy of ~13% (**Fig. 2i**), demonstrating the key role of competition between intramolecular interactions within the acceptor peptide with binding of regions of the acceptor peptide beyond the sequon to the extended acceptor binding surface in Stt3.

Pocket 2 regulates OST activation with a ‘switch’ mechanism, discussed below. It accommodates the residue at position −1 and the backbone of residue −2 of the acceptor substrate. For some peptides, binding at Pocket 2 is critical for their optimal orientation in the binding groove, such as the hydrogen bonding interaction between Glu45 of Pocket 2 and the Ser at −1 in PLB2_N195. In general, the structure and electrostatic profile of the residues in Pocket 2 allow for a wide variety of sidechains.

Pocket 3 accommodates residues at position +1 of the acceptor substrate, which can be any residue except Pro. The accessible volume within Pocket 3 is smaller than Pocket 1 and 2, but an *in silico* mutagenesis screen of residues at +1 on the equilibrated backbone structure of the model peptide YJR1_N99 showed that despite this limited accessibility, when the acceptor peptide substrate is optimally docked, Pocket 3 can accommodate a wide range of sidechains in terms of both steric and electrostatic considerations, because the sidechain of the residue at position −1 is directed outwards (**Fig. S4**). The limited volume of Pocket 3 likely contributes to constraining the dynamics of large sidechains.

To verify the role of the amino acids lining the substrate complementarity pockets in Stt3, we performed alanine-scanning mutagenesis of the residues lining Pocket 1 and 2 (**Fig. 2h**). To achieve this, we used a TetO7-*STT3* yeast strain in which expression of genomic *STT3* can be repressed by addition of doxycycline, while variant *STT3* can be expressed from a plasmid. As we predicted that mutations in Pocket 1 would affect the peptide-acceptor specificity of OST, we used LC-MS/MS to measure site-specific glycosylation occupancy in yeast expressing these Stt3 variants. We found that most *N*-glycosylation sites were efficiently glycosylated in cells expressing plasmid borne wild-type Stt3, while in the negative control strain treated with tunicamycin many *N*-glycosylation sites were inefficiently glycosylated. In contrast, yeast expressing Stt3 variants with mutations to Ala in Pocket 1 displayed subtle and distinct site-specific under-glycosylation phenotypes (**Fig. 2g** and **Fig. S7**), consistent with the amino acids lining these pockets in Stt3 allowing interactions with specific acceptor peptides through diverse interaction mechanisms.

The extended interface for interactions between diverse acceptor substrate peptides and OST we identify here is consistent with previous observations of the substrate specificities of eukaryotic and bacterial *N*-glycosylation machineries. In mammalian cells, “extended aromatic sequons” (EAS) with aromatic residues at the −2 or −3 position have been reported to be glycosylated with high efficiencies^15,22,23^. This high glycosylation efficiency is likely driven by interactions between the side chain of these aromatic residues and the hydrophobic Pocket 1 in Stt3. In the more distantly related bacterial *N*-glycosylation system, PglB, the bacterial homolog of Stt3, glycosylates Asn in an extended sequon D/E-x-N-x-S/T^6,20,24,25^. In PglB, this sequon is recognized and bound with high affinity through a salt bridge with an Arg in PglB, located at the equivalent position to the Pocket 1 in eukaryotic Stt3. This suggests that the extended interactions we observe between Stt3 and acceptor peptides beyond the sequon is an ancient evolutionary feature of *N*-glycosylation machineries.

### OST catalytic activity is turned on by a ‘switch’

Analysis of our MD simulations allowed us to identify a unique mechanism, which we name a ‘switch’, that activates OST catalysis by controlling the release of the assisting base Glu350. In the cryo-EM structure we used to build all our 3D model complexes (PDB 8AGC) OST is bound to a synthetic peptide mimetic inhibitor^19^ and corresponds to a catalytically inactive state, with Glu350 engaged in a salt bridge with Arg159 (**Fig. 3a)**. We define this resting state as ‘OFF’. Binding of an acceptor peptide substrate in a productive conformation, such as seen spontaneously during MD simulations of OST in complex with the YJR1_N99 and PLB2_N193 peptides, both of which are efficiently glycosylated, favours a structural rearrangement to the catalytically active alignment of the Asp47 nucleophile and Glu350, which is released from the salt bridge with Arg159. This conformational change is mechanistically triggered by optimal binding of the acceptor peptide at Pocket 1 or 2 positioning the carbonyl of the backbone of the acceptor peptide at −2 close to the carboxylate of Glu350 in the OFF state, and electrostatic repulsion triggering a switch to the ‘ON’ position, corresponding to the catalytically active state in which the Arg159 forms a salt bridge with Glu45, and Glu350 is free to act as the assisting base (**Fig. 3b** and **3c)**. Notably, site-directed mutagenesis of R159A in our TetO7-STT3 yeast system was lethal to yeast (**Fig. S6**), consistent with the essential role of Stt3 Arg159 in the regulation of OST activity.

**Figure 3.**
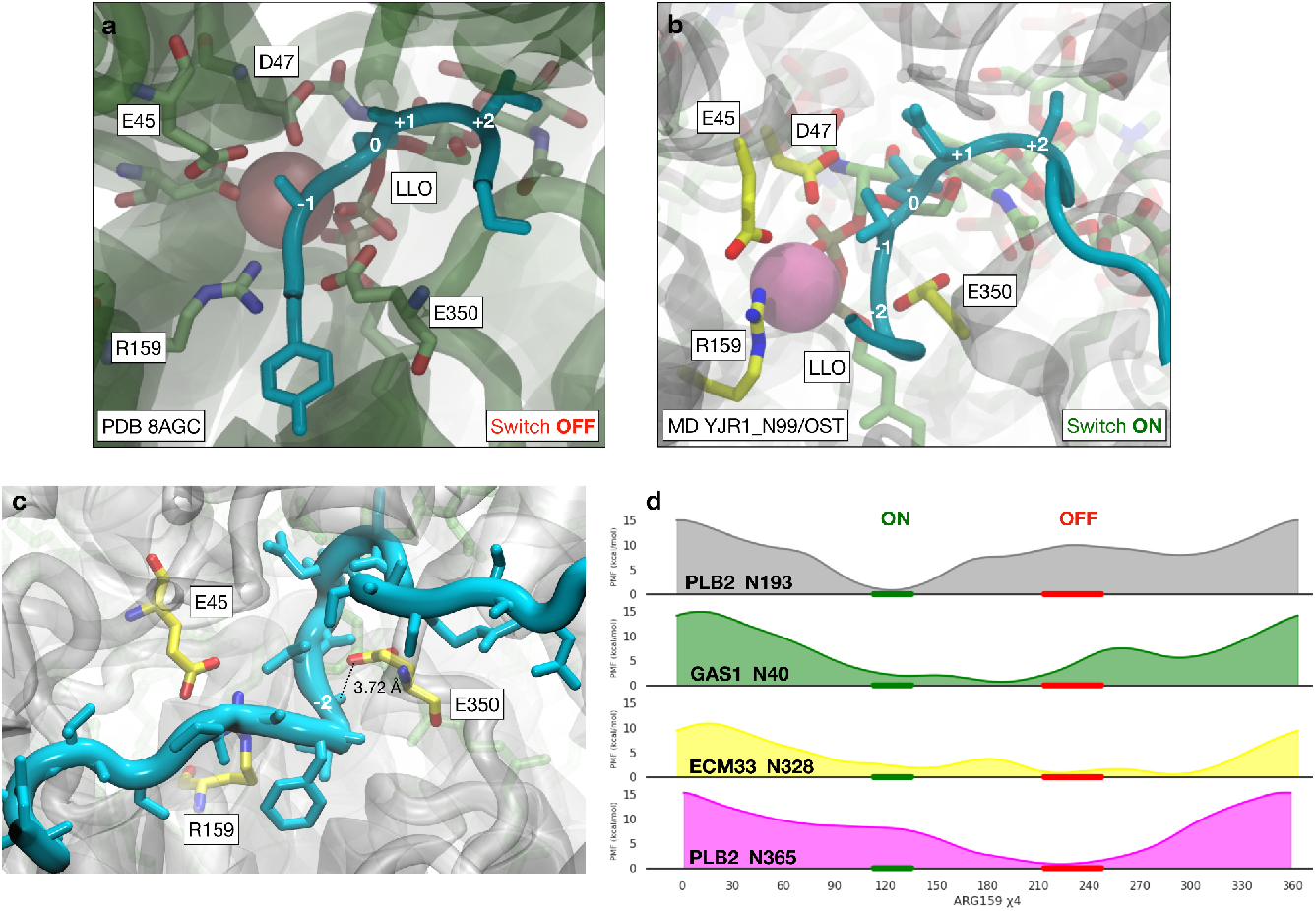
An ON/OFF switch in OST is controlled by high affinity acceptor peptide binding. **(a)** Close-up view of the OST structure in complex with a substrate mimetic rendered in cyan cartoons (PDB 8AGC). Labels indicate positions from −1 to +2 along the substrate. Key catalytic residues are labelled and rendered with sticks with C atoms in green, N atoms in blue and O atoms in red. In this structure the switch is defined as “OFF” as the catalytic base Glu350 is engaged in a salt bridge with Arg159. Asp47 is the catalytic nucleophile. **(b)** Representative snapshot from the equilibrated conformation of the complex between the OST model and the efficiently glycosylated YJR1_N99, corresponding to 520 ns of the 1 μs trajectory. The presence of an active substrate in the binding site with the Asn acceptor in a productive configuration displaces the Glu350 for the reaction to proceed, releasing Arg159 to form a salt bridge with Glu45, switching ‘ON’ the OST. **(c)** Close up view of the OST switch in the ON state from the snapshot from the OST/YJR1_N99 MD trajectory at 520 ns. The Mn^2+^ ion is not represented for clarity. The carbonyl of the residue at −2 triggers the opening of E350-R159 salt bridge in the resting OFF state by electrostatic repulsion with the carboxylate of E350, located 3.72 Å away in this frame. **(d)** Potential of mean force (PMF) rendered with spline regression, calculated for the OST in complex with PLB2_N193 (grey; efficiently glycosylated), GAS1_N40 (green) and ECM33_N328 (yellow) (both underglycosylated under stress), and PLB2_N365 (pink; never glycosylated). The labels ON (green) and OFF (red) indicate active and inactive orientations of the Arg159 sidechain, respectively, corresponding to values between 120°–145° (for ON), and 210°–240° (for OFF). Molecular structures rendered with Visual Molecular Dynamics^21^ (VMD) (https://www.ks.uiuc.edu/Research/vmd/). PMF data visualization rendered with *seaborn* FacetGrid (https://seaborn.pydata.org/).

Our unbiased MD simulations reveal an additional interplay between acceptor substrate binding at Pocket 1 and Pocket 2 with activation of the OST switch. These simulations showed that when the acceptor substrate is optimally docked through stable interactions across Pocket 1, 2, and 3, the sidechain of the residue at position −1 was directed away from Glu45, enabling a broad range of residues to occupy this position. However, when the acceptor substrate was not complementary to Pocket 1, the residues at position −1 could be re-orientated and interfere with the switch by displacing either Glu45 and/or Arg159. For instance, MD simulation of the complex with PLB2_N365 that was never glycosylated showed that the Asn364 at position −1 and Lys361 at position −4 both formed hydrogen bonds with Stt3 Glu45, hindering Arg159 from engaging with the assisting base and stabilizing the switch in the OFF position.

To understand the basis by which various substrate peptides could differentially activate the OST switch, and based on the analysis of the spontaneous activation of the switch we observed during the deterministic trajectory of OST in complex with YJR1_N99, we calculated the free energy involved in activating the switch with a range of different acceptor peptides bound. The potential of mean force (PMF) was determined using the rotation of the Arg159 χ_4_ torsion angle as the reaction coordinate for the complexes with PLB2_N193 (efficiently glycosylated), GAS1_N40 and ECM33_N328 (both underglycosylated under stress), and PLB2_N365 (never glycosylated). All PMF calculations started from an equilibrated state of the corresponding complex selected from the 1 μs unbiased MD simulations. The results indicated that the binding of efficiently glycosylated PLB2_N193 led to a free energy minimum corresponding to the ON position of the switch, while the OFF conformation corresponded to an increase in free energy of 10.3 kcal/mol (**Fig. 3d**). For the complexes of OST with peptides that were glycosylated in optimal conditions but underglycosylated under stress, GAS1_N40 and ECM33_N328, the PMF indicated that both ON and OFF states were energetically accessible at room temperature. Meanwhile, for the never glycosylated PLB2_N365 the free energy minimum corresponded to the OFF position of the switch, with an increase in free energy up to 8.2 kcal/mol for the ON state.

Together, these results show that only acceptor peptide substrates that bind to OST with high affinity can trigger the switch to turn ON, providing tight control of OST activity, while the mechanism of OFF/ON switch activation hinges on electrostatic repulsion exerted by the backbone carbonyl group of the −2 position of the acceptor peptide, explaining how OST activation is not determined by a simple substrate sequence or motif.

### Analysis of buried sequons reveals sequence motifs controlling *N*-glycosylation efficiency

The factors that determine whether a particular Asn is glycosylated or not are poorly understood. Our MD simulations of the yeast OST in complex with preferred and suboptimal peptide substrates showed how the interactions established with the extended binding groove regulate OST’s *N*-glycosylation efficiency for a broad variety of substrates. Optimal substrates are highly complementary to the extended groove, with acceptor peptide sidechains establishing interactions across Pockets 1-3 that lead to appropriate positioning of the backbone carbonyl of the −2 position and activation of the ON switch. Our results also showed that the loss of binding to Pocket 1 tended to destabilise the N-terminal side of the acceptor peptide from the optimal bound pose, leaving the switch in the OFF resting state, and decreasing *N-*glycosylation efficiency.

Based on these results, we inferred that the main requirement for an Asn to be able to be glycosylated by OST is for it to be located in a stretch of acceptor peptide substrate that can adopt a conformation complementary to the extended OST groove, allowing at least some degree of binding. The most important element of the acceptor peptide is the sequon itself. The N-x-S/T region of well-bound acceptor peptides is exceptionally extended, as shown in the conformation of the acceptor peptide substrate in a cryo-EM structure of OST with bound peptide substrate (PDB 8AGE), with φ=-154.42° and ψ=+156.91° at the +1 position (red dot in **Fig. 4a-f**). These structure requirements are ultimately the reason why Pro cannot occupy the +1 position in the N-x-S/T sequon, because its presence constrains the backbone in a conformation incompatible with the binding requirements of the OST catalytic site^5^. We confirmed this mechanism by calculating the Ramachandran plots for all N-P-S/T sequences in all human proteins in the PDB (total of 15,568 sequences in 2,815 proteins without duplicates) with resolution ≤ 3.0 Å. These results clearly indicated that the local conformation of the backbone corresponding to a sequon with Pro at +1 strongly deviated from the optimal glycosylation-competent conformation of peptide bound to OST (**Fig. 4a, b**, and **g)**. We extended this analysis to include all other N-x-S/T sequons in this data set, which identified a wide range of complementarity. Optimal consistency with OST binding mainly depended on the identity of the x amino acid at +1 (**Fig. 4g**), but also with a structure-based preference for Thr over Ser at +2, with 70% of the 20 sequons with the highest structural complementarity carrying a Thr at +2 (**Fig. 4g)**. We augmented this structural analysis by considering all (human and other) proteins from UniRef50 with the corresponding structures predicted by AlphaFold 2^26,27^ filtered for sequons where the *per-*residue confidence score (pLDDT) was ≥ 90 across the full −4 to +4 window, for a total of 1.82 million sequons in 1.00 million proteins (**Fig. S9, S10**, and **S11**). In summary, these analyses showed that the propensity for the N-x-S/T sequon backbone to adopt an extended conformation compatible with optimal OST binding explains a key mechanism by which the amino acid residue at the +1 position strongly affects glycosylation efficiency^8,13,14,28^.

**Figure 4.**
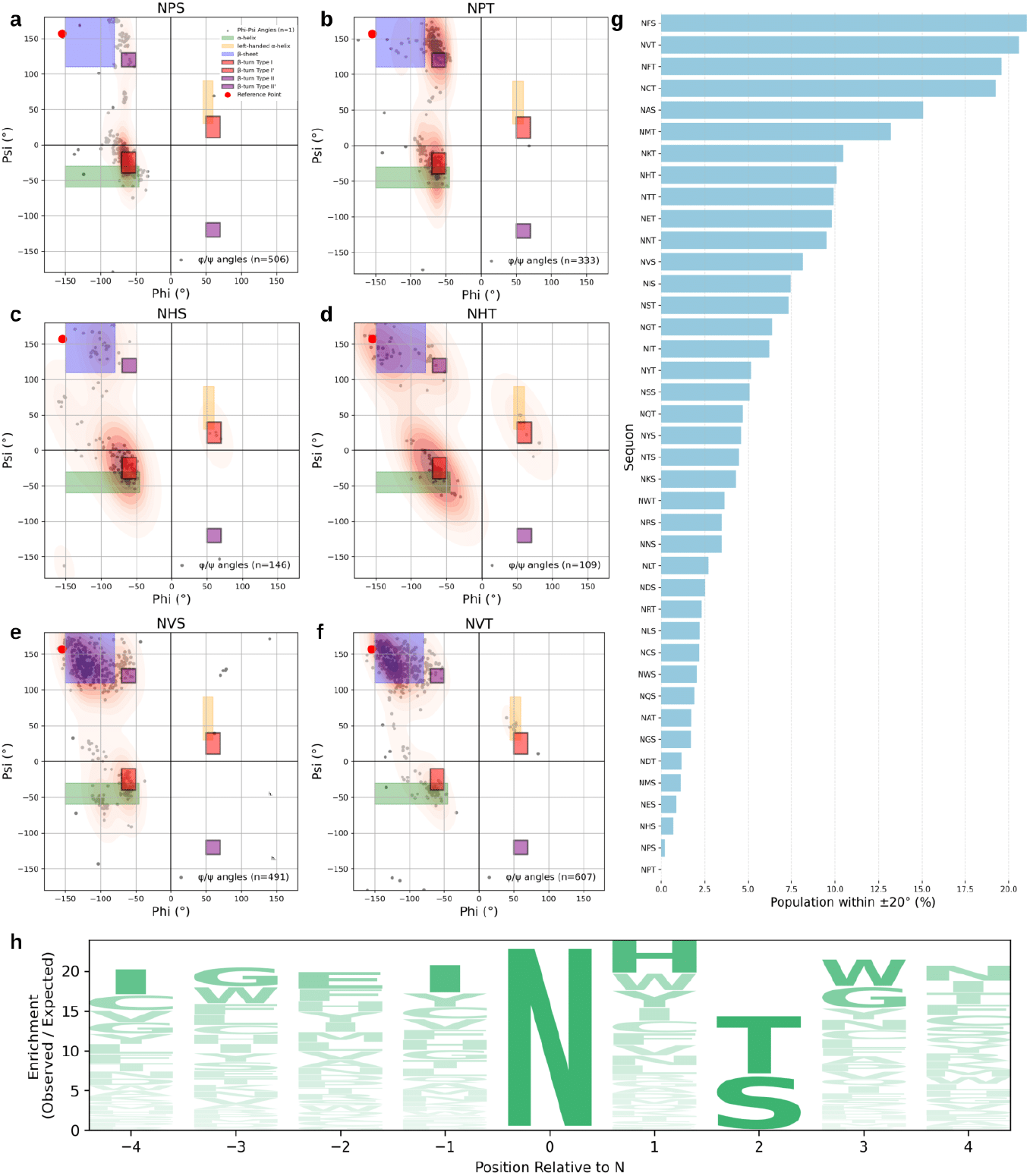
Features of surface exposed and buried sequons. Ramachandran plots of φ/ψ angles at position +1 for selected canonical sequon subtypes: **(a)** N-P-S, **(b)** N-P-T, **(c)** N-H-S, **(d)** N-H-T, **(e)** N-V-S, **(f)** N-V-T. Data points correspond to experimentally resolved human protein structures in the PDB with resolution ≤ 3.0 Å. Each heat map shows the distribution of dihedral angles (gray dots) and kernel density estimates (red contours). Colored boxes indicate regions associated with secondary structures: α-helix (green), β-sheet (blue), left-handed α-helix (yellow), and β-turn types I, II, I’, II’ (purple). The red dot marks the φ/ψ region of the glycosylated acceptor peptide bound to yeast OST (PDB: 8AGE), used as a reference for catalytically competent conformations. **(g)** Bar plot indicating the relative proportion (%) of sequons in all secreted proteins in the PDB with φ/ψ conformation within +/-20^□^ of the reference structure. **(h)** Amino acid enrichment across the –4 to +4 region surrounding buried canonical sequons (internal sequons). Letter height reflects the ratio between observed and expected frequencies based on vertebrate background distributions. Enrichment of turn-promoting residues such as His and Trp at position +1 supports a conformational mechanism that disfavors glycosylation. Plots were generated using *Matplotlib* v3.8.4.

A critical aspect of the regulation of OST is that Asn that are buried in the folded structure of a protein must not be glycosylated, as glycosylation at these sites would be incompatible with productive folding of the protein. We therefore suspected that Asn in sequons located in the interior of secreted proteins would have local sequence contexts that reduce OST activity. To identify these sequences we extracted canonical *N*-glycosylation sequons, (N-x-S/T; x ≠ P) from the reviewed and complete set of secreted human proteins available in UniProtKB^29^, resulting in 14,835 sequons from 3,648 proteins. To filter for internal sequons we evaluated the local structural accessibility by calculating the relative solvent accessibility (RSA) of the Asn using the DSSP module in Biopython^30^. The RSA value provides a normalized measure of the residue’s exposure to the solvent. For each protein we calculated RSA values from the corresponding 3D structures in the PDB where available (541 proteins) or high-confidence (pLDDT ≥ 90) predicted models from the AlphaFold Protein Structure Database^27^ (257 proteins). We classified sequons as ‘buried’ when the corresponding Asn RSA value was below a threshold of 15%, a value slightly more stringent than the default value of 16%^31^. The results of this analysis identified an enrichment of specific residues surrounding the sequon (**Fig. 4i)**, with the sequence IGEI<u>N</u>H[S/T][W/G]N based on the analysis of 997 sequons representing an inherently non-preferred acceptor substrate for OST.

The enrichment of His and Trp at +1 is a striking feature of the internal, non-glycosylated sequons (**Fig. 4h**) that is consistent with our analysis of the secondary structure of all N-x-S/T sequons from human proteins in the PDB (**Fig. 4g**). In particular, N-**H**-S sequons tend to adopt a secondary structure incompatible with OST binding (**Fig. 4c** and **g**), almost to the extent of the completely non-glycosylatable N-**P**-S/T sequences. On the other hand, N-**H**-T sequons are more compatible with OST binding (**Fig. 4d** and **g**), demonstrating that the presence of Thr at the +2 position can act to increase glycosylation directly through higher complementarity to the OST, as well as through indirect effects on substrate secondary structure. Other sequons (**Fig. 4h**), especially those with small hydrophobic amino acids such as N-**V**-S (**Fig. 4e**) or N-**V**-T (**Fig. 4f**) and others (see **Fig. S4**), can more readily access regions corresponding to the OST bound peptide (PDB 8AGC) consistent with these residues not being enriched in internal sequons (**Fig. 4i)**. Other features of this representative non-preferred OST substrate are also noteworthy. The results of our MD simulations showed that Gly at position −3 does not effectively engage with Pocket 1, and that Glu at position −2 could interfere with residues Arg159 and Glu45 that control the OST ON/OFF switch. Other enriched features in this representative non-preferred substrate are more difficult to interpret outside of a specific sequence context, which is possibly why enrichment at some sites is more subtle. The absence of strong determinants of poor glycosylation across the full −4/+4 window also allows OST to maintain broad sequence specificity.

### Sequons that are poorly accommodated by OST cause low *N*-glycan occupancy in human glycoproteins

A common challenge in heterologous protein expression in mammalian cells is the difficulty in predicting whether an Asn in a glycosylation sequon will be efficiently glycosylated or not. For example, Fab FL14 derived from a patient with follicular lymphoma contains two *N*-glycosylation sequons, Asn38 and Asn57, but LC-MS/MS analysis shows that only Asn38 is glycosylated^32^. This difference in glycosylation occupancy is not due to structural constraints, as the crystal structure of Fab FL14 shows a solvent-exposed but unoccupied Asn57 while glycosylation of the nearby Asn38 is strongly resolved in the density (**Fig. 5b**). This suggested that the lack of glycosylation at Asn57 was due to its location within a sequence context that was not a preferred substrate of OST.

**Figure 5.**
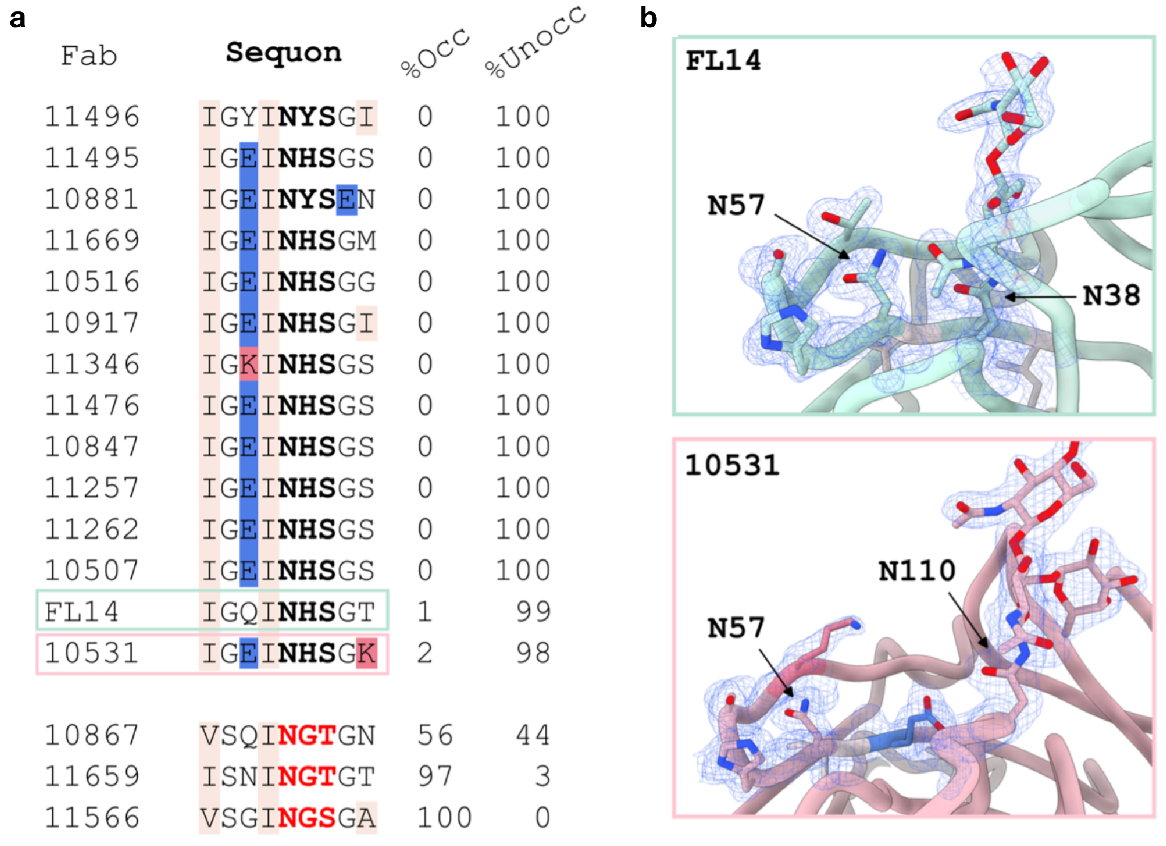
Local sequence context controls glycosylation occupancy at Asn57 in human Fabs. **(a)** Percentage occupied (%Occ) and un-occupied (%Unocc) by *N*-glycosylation at Asn57 for each Fab, as determined by LC-MS/MS. Extended sequence context of Fabs centered at Asn57 with the classical N-x-S/T sequon in bold. Residues with negative charge (glutamate) are blue, positive charge (lysine) are pink, and aliphatic residues (isoleucine, valine and alanine) are beige. **(b)** Crystal structures of previously reported FL14 Fab (PDB *submission pending*) and 10531 Fab resolved to 1.76 Å, containing solvent-exposed, but unoccupied, Asn57. The final 2*F*_obs_-*F*_calc_ electron density maps (contoured at 1.1 sigma) are shown for the extended Asn57 sequons and coloured as in (a). Density is also shown for occupied glycosylation sites in each Fab (Asn38 for FL14, and Asn110 for 10531).

Intriguingly, the sequence context of the non-glycosylated N57 in Fab FL14 is IGQI**N**HSGT, which bears a remarkable similarity to the representative sequence for inherently non-preferred acceptor substrates for OST, IGEI**N**H[S/T][W/G]N (**Fig. 4h**), discussed above. To more closely validate the link between unfavourable acceptor peptide-OST interactions and poor glycan occupancy, we expressed an additional 16 Fabs containing the N57 site in various local sequence contexts in HEK293F cells^33^. We solved the crystal structure of a representative Fab 10531, which again showed clear electron density for the region surrounding Asn57 (IGEI**N**HSGK), but with no density consistent with a *N*-glycan at this site. Asn57 has a solvent-accessible surface area of 23.2 Å^2^ as calculated by PDBePISA^34^ and thus, based on the 3D structure alone, would be able to accommodate an *N*-glycan. We then analyzed the glycan occupancy of these 16 Fabs by LC-MS/MS. This analysis revealed persistently inefficient glycosylation at Asn57 with a series of amino acid changes introducing Glu, Lys, or Tyr at −2, while leaving unchanged residues important for the local structural context (<u>IG</u>x<u>I</u>**N**) (**Fig. 5a**). The lack of glycosylation with these sequences is consistent with the mechanisms underlying efficient glycosylation by OST described above: Glu or Lys at −2 can interfere with the OST residues controlling the ON/OFF switch (Arg159 and Glu45 in Stt3); while Gly at −3 or Tyr at −2 do not contribute to interactions with Pocket 1, leaving the N-terminal acceptor peptide tail unbound. We next considered Fabs with different amino acids at the +1 position, as we had identified N-**H**-S sequons as particularly poor substrates for OST (**Fig. 4**). Fabs with the His at +1 replaced by the structurally conservative Tyr did not increase glycosylation occupancy (**Fig. 5a**), consistent with the lack of enrichment of N-Y-S sequons in buried sequons (**Fig. 4h**), but introduction of Gly at +1 strongly increased glycosylation occupancy in a variety of sequence contexts, particularly in Fab 11566, where the sequence VSGIN**G**SGA also lacks the nonideal His at +1, Glu at −2, and Gly at −3 (**Fig. 5a**). In summary, the dramatic changes in glycan occupancy at this site despite conservation of its structural context confirm that *N*-glycosylation efficiency is dictated by the extended sequence context beyond the classic N-x-S/T motif.

## Conclusions

Our combined analyses have characterised the molecular determinants that regulate the substrate specificity and catalytic efficiency of eukaryotic OST. We find that to be efficiently glycosylated, an Asn must be: located in an N-x-S/T sequon; in a stretch of polypeptide substrate without secondary structure, including an amino acid residue at the +1 position with an appropriately extended Asn-Xaa peptide bond conformation consistent with optimal interaction with OST; in a sequence context with low propensity to form a hairpin, either through hydrophobic or salt bridge interactions; and with a local sequence that can tightly interact with Pockets 1, 2, and 3 to activate the ON switch in OST. Within this framework, this in-depth characterization of the molecular mechanisms regulating OST catalytic efficiency will increase our understanding of the constraints of secretory protein sequence and evolution, enable the prediction of changes in viral antigenicity and immunity^35–37^, allow prediction of how genetic mutations affect glycosylation as a trigger for pathogenic phenotypes^38–41^, and assist the design of biologics and other designer glycoproteins with desired site-specific glycosylation.

## Acknowledgements

We acknowledge the technical assistance and expertise of The University of Queensland, School of Chemistry and Molecular Biosciences Mass Spectrometry Facility. We gratefully acknowledge the Science Foundation of Ireland (SFI) Centre for Research Training in Foundations of Data Science (www.data-science.ie) for financial support of BT postgraduate training (18/CRT/6049), National Health and Medical Research Council (NHMRC) Ideas grant APP1186699 to BLS, and Australian Research Council (ARC) Linkage Infrastructure, Equipment and Facilities grant LE220100068 to BLS.

