## Supplementary Material for "Regulation of *N*-glycosylation efficiency by eukaryotic oligosaccharyltransferase"

### Results

**Titration of glycosylation stress conditions.** We studied the differential glycosylation activity of OST with suboptimal LLO donor substrate (Man9GlcNAc2), and mature LLO substrate (Glc3Man9GlcNAc2) at low LLO concentration. We compared site-specific glycosylation in a yeast strain lacking the gene encoding the Alg6 glucosyltransferase (Δ*alg6*), to a yeast strain deficient in Alg7 activity, which we created through chemically-treating wild type cells with the antibiotic tunicamycin (Tm). We expected each strain would produce phenotypically distinct defects, whereby the Δ*alg6* mutant would produce the truncated Man9GlcNAc2 LLO at normal cellular levels, whereas the alg7 deficient strain would produce the full-length LLO (Glc3Man9GlcNAc2) at reduced cellular levels. In order to directly compare site specific glycan occupancy between these strains, we created similar global hypoglycosylation phenotypes between the Δ*alg6* cells and Tm-treated cells. We first identified the concentration of Tm that would provide a similar overall under-glycosylation phenotype as the Δ*alg6* strain. We performed a titration experiment that tested a series of concentrations of Tm. We grew Δ*alg6* cells, and wild type BY4741 supplemented with varying concentrations of Tm. We also used a separate system to generate glycan structural and concentration stress. In this system, we used the Tet-Off system, where the gene of interest (*ALG7* or *ALG6*) was brought under the control of Tet-R promoter and operator and the cells were treated with appropriate concentrations of the tetracycline analogue Doxycycline (Dox) to selectively down-regulate gene expression. We used the wild-type R1158 and Tet-R-*ALG7* strains from the Tet Hughes Collection (yTHC)[1]. The Tet-R-*ALG6* strain was created by insertion of the Tet-R promoter upstream of the *ALG6* gene via PCR amplification and homologous recombination. Previous studies found that Dox at concentration of 40 µg/mL is sufficient to completely inhibit the expression of a gene under the control of the Tet-R system and does not have any substantial effect on cellular morphology or global gene expression[2]. Therefore, we treated the Tet-R-*ALG6* strain with

Dox at 40 µg/mL concentration to completely inhibit the expression of *ALG6* to generate Alg6 deficiency. We grew Tet-R-*ALG7* cells in the presence of varying concentrations of Dox. After harvesting cells at mid-log phase, we measured site-specific *N*-glycosylation occupancy in yeast cell wall proteins[3,4]. We extracted yeast cell wall proteins, and then reduced, alkylated and digested proteins with trypsin, and employed SWATH-MS to measure peptide abundance and site-specific hypoglycosylation at each Tm concentration compared to Δ*alg6* (**Fig. S1a**) and at each Dox concentration for Tet-R-*ALG7* cells compared to Tet-R-*ALG6* cells grown with 40 µg/mL Dox (**Fig. S1b**). Although there was high variability in the glycan occupancy measured at different sites, the average occupancy in yeast cells supplemented with 0.14 µg/ml of Tm was similar to the Δ*alg6* cells, while the average occupancy in Tet-R-*ALG7* cells with 0.1 µg/mL was similar to Tet-R-*ALG6* cells grown with 40 µg/mL Dox. We therefore used these concentrations for comparison of site-specific glycosylation occupancy in our screen (**Fig. S2**).

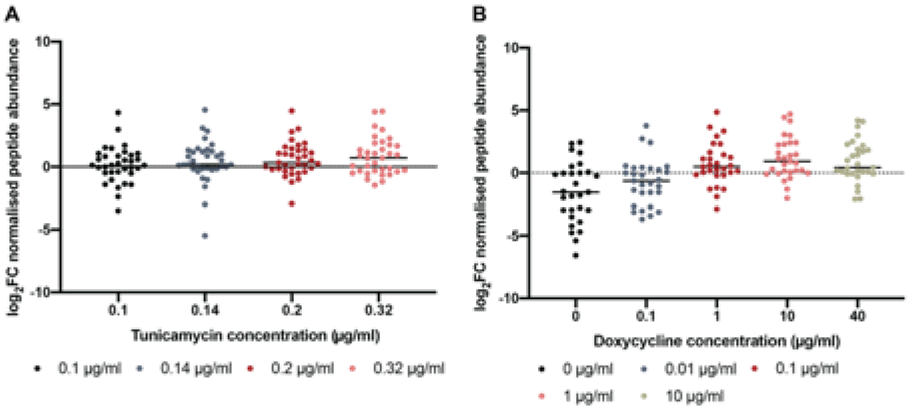


**Figure S1. Tunicamycin (Tm) and doxycycline (Dox) concentration titration assays. (a)** Average log2FC normalised sequon-containing peptide abundance of cell wall proteins in BY4741 yeast treated with various concentrations of Tm compared to the Δ*alg6* strain. **(b)** Average log2FC normalised peptide abundance in the Tet-R-*ALG7* strain treated with various concentrations of Dox compared to the Tet-R-*ALG6* strain treated with 40 µg/mL of Dox. Horizontal lines indicate average log2FC of normalised peptide abundance with Alg7 deficiency compared to Alg6 deficiency.

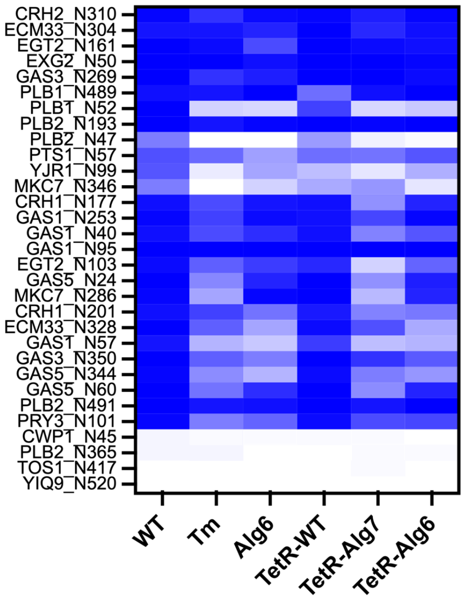


**Figure S2. *N*-glycosylation site occupancy in wild-type yeast and yeast subject to glycan structural or concentration stress.** WT, wild type BY4741; Tm, BY4741 treated with 0.14 µg/mL tunicamycin; Alg6, Δ*alg6* yeast; TetR-WT, wild type R1158; TetR-Alg7, TetR-*ALG7* treated with 0.1 µg/mL Dox; TetR-Alg6, TetR-*ALG6* treated with 40 µg/mL Dox. Each square in the heat map represents the mean of biological replicates (n=6) coloured from white (0% glycosylated) to blue (100% glycosylated).

To analyse the impact of Asp44 in Cwp1 on glycosylation of Asn45, we cloned Cwp1 and variant Cwp1_D44V, both with a C-terminal His-tag, and expressed the proteins in wild type yeast. We purified the Cwp1 proteins from the supernatant, reduced/alkylated cysteines, deglycosylated with EndoH, digested with trypsin, and measured peptides and deglycosylated peptides with LC-MS/MS. We could identify only non-glycosylated Asn45 from Cwp1, but both glycosylated and non-glycosylated Asn45 from Cwp1_D44V (**Fig. S3**).

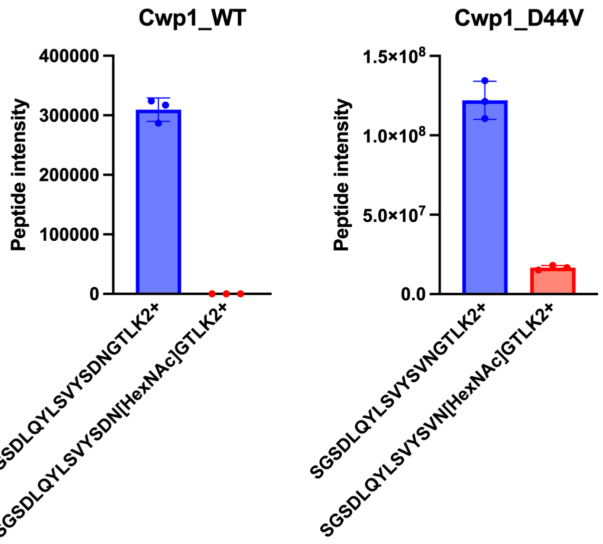


**Figure S3. *N*-glycosylation site occupancy at Asn 45 in wild type Cwp1 and Cwp1_D44V purified from yeast.** Cwp1 and Cwp1_D44V with C-terminal His-tags were expressed and purified from wild type yeast, digested with EndoH and trypsin, and measured with LC-MS/MS. Graphs show the intensity of non-glycosylated and previously N-glycosylated peptides. Data points show values from triplicate analyses. Occupancy is shown in Fig. 2i.

**Single-point *in silico* scanning of the substrate peptide at position +1.** Our conformational analysis based on the MD simulations of six OST-peptide complexes suggested that the interactions that peptide substrates form along the OST extended binding site are crucial to modulating its affinity, thus favouring or impeding catalytic activity. Based on these results we identified three regions along the OST binding groove, named Pocket 1, 2 and 3, that are critical for supporting a productive conformation of the substrate peptide between a -4 to +4 stretch. Pocket 1 hosts the sidechains of residues at -3 (or possibly at -2), pocket 2 hosts the residue at -1 and the OST switch residues, namely Glu45, Arg159 and the catalytic assisting base Glu350, while Pocket 3 is the narrowest region, framed by Phe46, Tyr 519 and Asp518 accommodating the sidechain of the residue at +1 in the sequon. Based on the equilibrated structure of the complex with the YJR1_N99, which has an NNS sequon, we ran a scan through all the possible 18 residues to evaluate the steric hindrance corresponding to different sidechains at this site. Selected representative cases are shown in **Fig. S4**. The scanning was done with the *Mutagenesis* tool in pymol (www.pymol.org) by selecting the allowed conformation of the sidechain corresponding to the minimum steric hindrance.

The results are qualitative, as the optimal accommodation of the sidechain at +1 depends on the substrate context in which the sequon is found. Nonetheless, within this purely structure-based approximation, Pocket 3 can accommodate virtually any sidechain (**Fig. S4)**. The amphipathic nature of the pocket, with Phe46 and Tyr 519 on one side and Asp518 on the other, also permits a broad selection of chemical profiles, from hydrophobic sidechains to hydrophilic and amphipathic. It is also noteworthy that the confinement of the sidechain at +1 within a narrow pocket while chemically supported by interactions with Pocket 3 residues, contributes to restraining the peptide’s dynamics, favouring efficient catalysis.

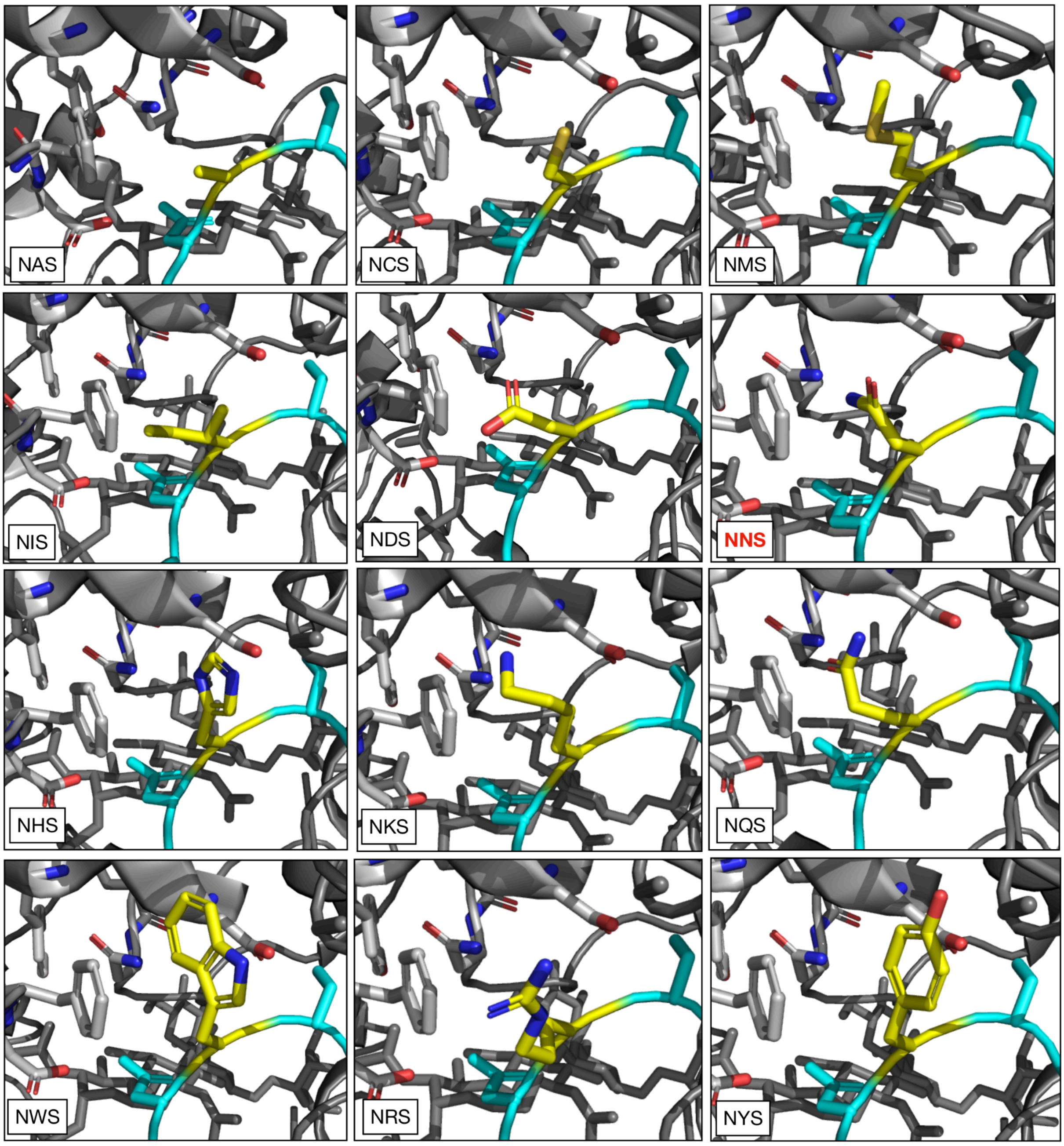


**Figure S4. Structures obtained from the *in silico* mutagenesis screening of residues at position +1** (yellow C atoms) run on a representative conformation (sampled at 520 ns) of the YJR1_N99 peptide backbone (cyan C atoms) from the MD simulation of the complex. The YJR1_N99 sequon (NNS) is highlighted with a red label. The key residues lining Pocket 3 are shown with light grey sticks, namely Phe46 and Asp518, shown in each frame on the left and right of the +1 sidechain, respectively. Tyr 519 is at the top of each +1 sidechain, but not captured in any of the frames. The remainder of the protein is rendered with grey cartoons. Labels at the bottom left of each panel indicate the sequon represented, with the label in red corresponding to the native YJR1_N99 sequon. Scanning was done with the *Mutagenesis* tool in pyMol (www.pymol.org) where the sidechain conformations were selected based on minimum steric hindrance. Rendering with Visual Molecular Dynamics[5] (VMD) (https://www.ks.uiuc.edu/Research/vmd/).

**Tyr at position -3 cannot always be accommodated in Pocket 1.** Pocket 1 is a crucial region in the OST extended binding site, which accommodates the sidechain of residues at -3 or -2**.** As described in the main text, Pocket 1 is structured as a cavity lined by hydrophobic/aromatic sidechains with the rim bearing short hydrophilic sidechains. This chemical and structural arrangement is ideal for peptides bearing at -3/-2 either short hydrophilic residues or large aromatics such as Trp, or Phe, but not Tyr, whose phenolic sidechain is large and amphipathic.

In GAS1_40 the Tyr at –3 is accommodated in Pocket 1 where it establishes interactions with residues of Ost3 (Phe263, Gln262, Met406) (**Fig. S5**). This peptide represents a very particular case, where the Phe residues at –4 and –5 stabilize a productive conformation through stacking interactions with Stt3 Phe356, Phe359, and Phe496. In PLB2_N365 the Tyr at –3 is displaced towards Asp369 at position +4, forming a hydrogen bond (3.49 Å) that disrupts its ability to occupy pocket 1 (**Fig. S5**). This intrapeptide interaction represents a conformational constraint, affecting the optimal engagement with the catalytic site. In CWP1_N45, which is never glycosylated, the Asp at –1 forms a salt bridge interaction with Lys at +4 (2.68 Å) that could affect the productive conformation of the peptide. As a result pocket 1 remains unoccupied, further highlighting how unfavorable intrapeptide contacts can preclude efficient OST binding. As discussed in the main text, in support of this interpretation the mutation of Asp44 to Val, which removes the potential to form a salt bridge, increases OST activity and glycosylation at Asn45 (**Fig. 2i** and **Fig. S5**).

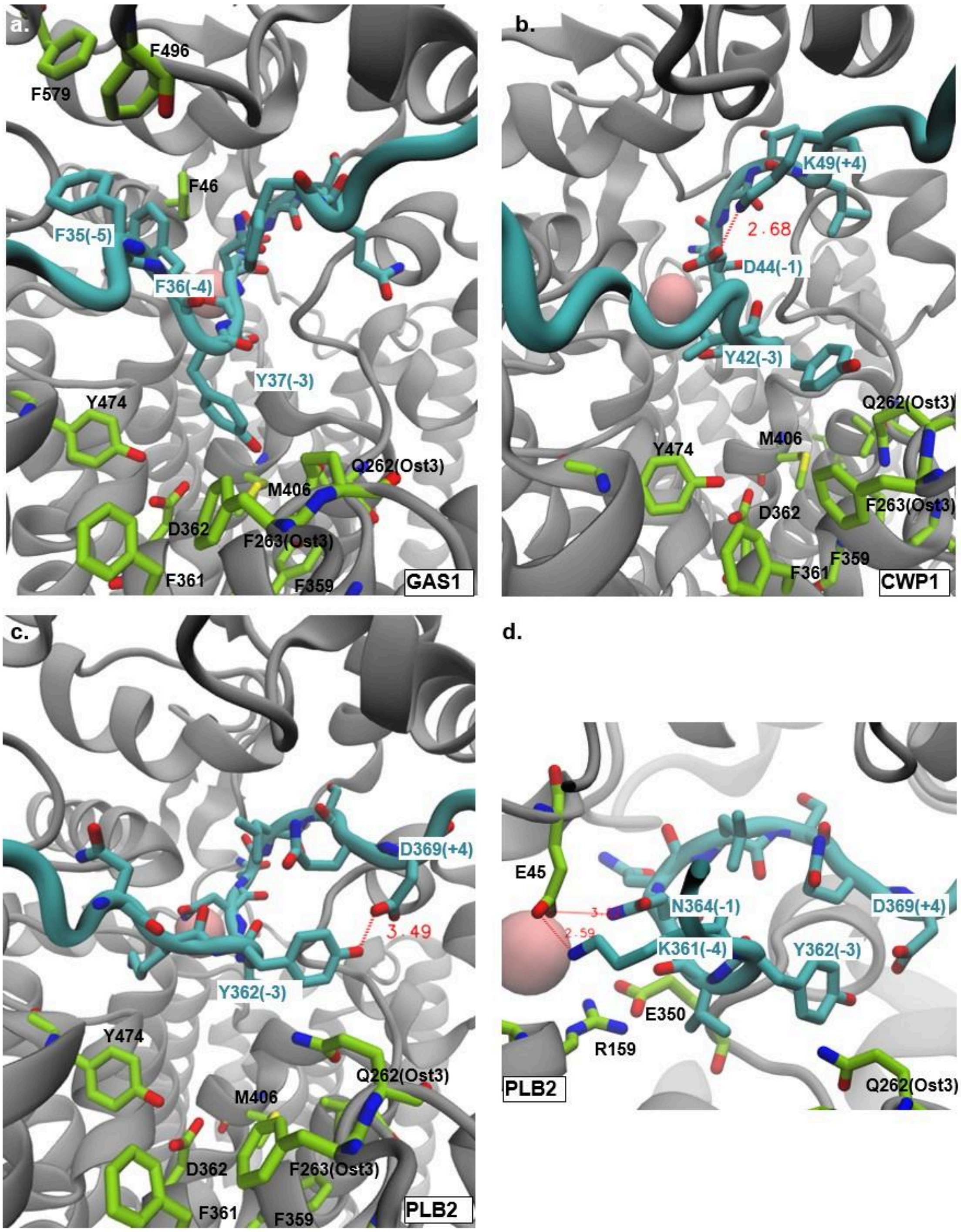


**Figure S5. Intrapeptide interactions modulate accessibility of pocket 1 in the OST binding groove.** Representative views of **(a)** GAS1_N40, **(b)** CWP1 and **(c)** PLB2_N365 peptides (cyan) in the binding site of OST. Key interacting residues from Ost3 are shown in green, while intra-peptide contacts are highlighted as red dashed lines with measured distances. GAS1_N40 displays optimal accommodation of the –3 Tyr in pocket 1 stabilized by surrounding aromatics, whereas in CWP1_N45 and PLB2_N365, unfavorable intrapeptide interactions at –1/+4 and –3/+4 prevent proper engagement of pocket 1. **(d)** Close up view of PLB2_N365: Asn364 at position -1 and Lys361 at position -4 both formed hydrogen bonds with Stt3 Glu45, hindering Arg159 from engaging with the assisting base E350, and stabilizing the switch in the OFF position. All structural images were generated with VMD (Visual Molecular Dynamics).

**Ala scanning to investigate the roles of residues in Pocket 1 and Pocket 2 of OST.** Stt3 is an essential subunit of the OST. As *N*-glycosylation is an essential modification in eukaryotes, complete deletion of an essential subunit of OST leads to cell death. Therefore, we used TetO7-*STT3* yeast, where genomic *STT3* is brought under the control of the Tet-OFF system. Treating TetO7-*STT3* yeast with doxycycline at the concentration of 10 µg/ml completely inhibited expression of genomic *STT3* resulting in no cell growth. We separately introduced the desired mutations in *STT3* expressed from the pRS413_STT3 plasmid. The plasmids pRS413, pRS413_STT3, or pRS413_STT3 with various mutations were transformed into the TetO7-*STT3* strain. Growing these transformed strains with 10 µg/mL Dox fully repressed expression of genomic STT3, such that the only potentially functional Stt3 is that from the pRS413 plasmid. We grew these strains on SD-His media with and without 10 µg/mL Dox. We found that the TetO7-STT3 pRS413 strain was not able to grow in the presence of Dox, as the strain could only express genomic *STT3*, transcription of which was under the control of Tet-OFF system and was inhibited due to the presence of Dox (**Fig. S6**). Strain TetO7-STT3 pRS413_STT3_R159A was also not able to survive in the presence of Dox, consistent with an essential role for Arg159 in the OST ON/OFF switch (see main text and **Fig. 3**). The remainder of the seven *STT3* variant strains grew equivalently to yeast expressing wild type *STT3* (**Fig. S6**). For the *STT3* variants which allowed yeast survival, we grew these yeast strains to mid-log phase, prepared cell wall fractions, and measured glycosylation occupancy with LC-MS/MS (**Fig. S7**).

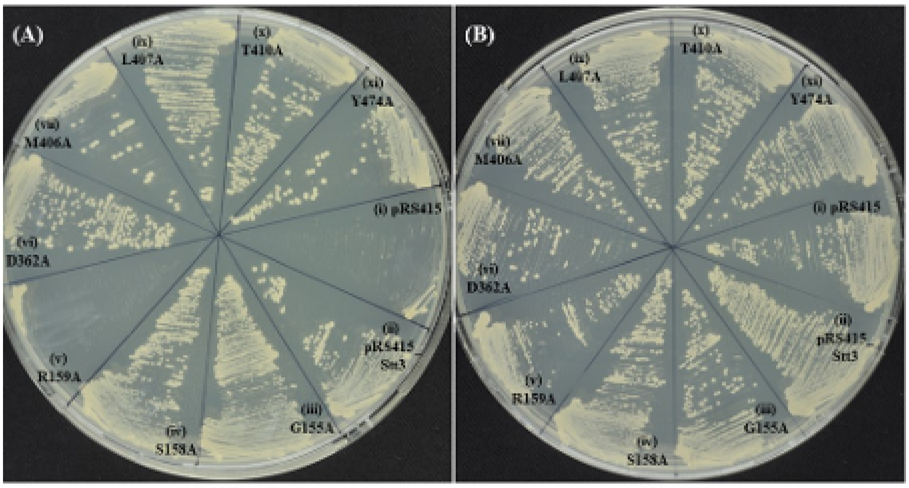


**Figure S6. Growth of yeast with variant *STT3* on SD-His media in the (A) presence and (B) absence of Dox.** (i) TetO7-STT3 pRS413 (ii) TetO7-STT3 pRS413_STT3 (iii) TetO7-STT3 pRS413_STT3_G158A (iv) TetO7-STT3 pRS413_STT3_ S158A (v) TetO7-STT3 pRS413_STT3_R159A (vi) TetO7-STT3 pRS413_STT3D362A (vii) TetO7-STT3 pRS413_STT3_M406A (viii) TetO7-STT3 pRS413_STT3_L407A (ix) TetO7-STT3 pRS413_STT3_T410A (x) TetO7-STT3 pRS413_STT3_Y474A.

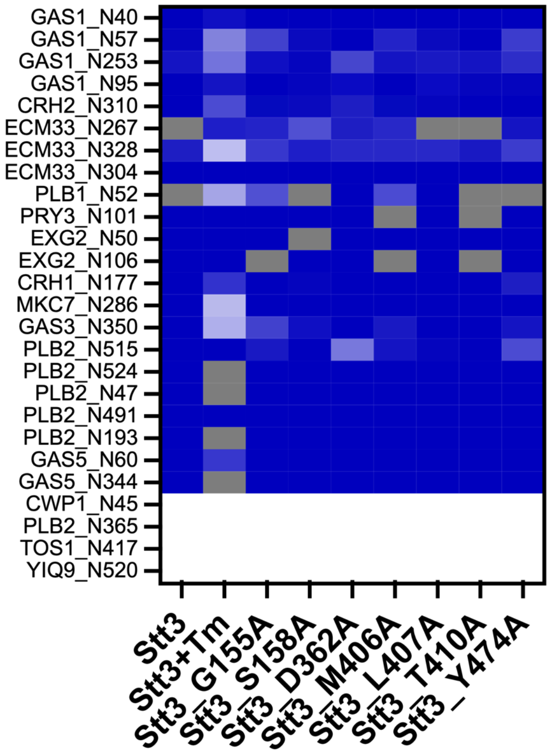


**Figure S7. *N*-glycosylation site occupancy in yeast expressing Stt3 Ala-scanning variants through Pocket 1 and 2.** TetO7-*STT3* yeast with expression of genomic *STT3* repressed with doxycycline and expressing plasmid-borne variant *STT3*. Stt3, native *STT3*; *Stt3*+Tm, native *STT3* plus 0.1 µg/mL tunicamycin; Stt3 variants with single Ala scanning mutations. Each square in the heat map represents the mean of biological replicates (n=6) coloured from white (0% glycosylated) to blue (100% glycosylated); grey, not measured. Fold change in site-specific glycosylation occupancy is shown in Fig. 2g.

**Substrate peptide dynamics: RMSD and RMSF Analysis.** The average RMSD values of the peptide's backbone atoms calculated along the MD trajectories are shown in **Table S1**. From these values it is clear that the dynamics of the peptides is largely at the N- and C-terminal tails, i.e. positions -5 to -10 and +5 to +10, respectively. These are not in contact with the OST. Meanwhile the section of the peptide substrate enclosed within the OST extended binding groove is dynamically restrained. Notably, only the Cα of the target Asn is restrained in place to avoid unphysical detachment or displacement of the peptide, yet the dynamics along the substrate is greatly reduced by interactions with the binding groove residues.

**Table S1. Root-mean-square deviation (RMSD) values (Å) of peptide backbone (BB) atoms calculated during the MD trajectories (1 μs)**. The backbone atoms (Cα,N,C) of the OST model were kept restrained throughout the MD trajectory, allowing only minor fluctuations of their relative positions. The full length of each peptide in the complex spans positions -10/+10 around the target Asn. The stretch within -4 and +4 aa encompasses residues in contact with the OST extended binding groove.

| Peptide | RMSD BB OST | RMSD BB peptide | RMSD BB peptide -4/+4 |
|---|---|---|---|
| YJR1_N99 | 0.34 | 3.71 | 1.24 |
| PLB2_N193 | 0.36 | 5.23 | 1.09 |
| GAS1_N40 | 0.34 | 3.29 | 1.07 |
| ECM33_N328 | 0.35 | 3.17 | 0.59 |
| PLB2_N365 | 0.34 | 2.77 | 0.97 |
| CWP1_N45 | 0.35 | 4.07 | 1.45 |

The RMSF values calculated for the residues along the full length of the peptide substrates show clearly that the highest flexibility is at the N- and C-terminal tails unrestrained by interactions between the peptide and the OST extended binding groove (positions -10 to -5 at the N-terminus and +5 to +10 at the C-terminus) (**Fig. S8**). The relatively large fluctuation of the YJR1_N99 Phe at -3 indicates the rotation of the Phe sidechain from outside to inside Pocket 1, where it rests for the majority of the 1 μs trajectory.

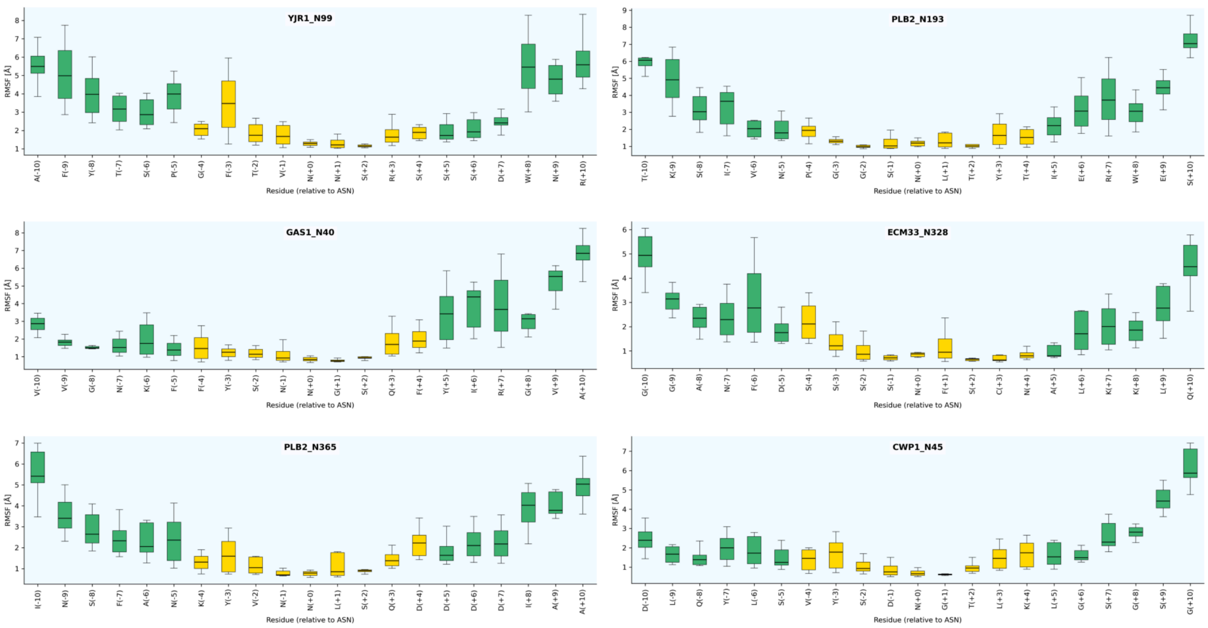


**Figure S8. Root-mean-square fluctuations (RMSF, Å) values calculated the residues along the full length (-10/+10 aa) peptide substrates.** RMSF values distributions are shown as boxplots for each residue, with positions indicated relative to the target Asn (position 0). Residues belonging to the –4 to +4 window are highlighted in yellow, while flanking residues are shown in green. Plots were generated with matplotlib and seaborn python libraries.

**Secondary Structure Analysis.** One of the fundamental determinants for *N*-glycosylation efficiency is sequence complementarity between the peptide substrate and the OST extended binding site. Local secondary structure features may prevent key interactions to occur and could determinants for poor *N*-glycosylation efficiency. Incompatible secondary structure motifs may be particularly disruptive around the sequon, where an optimal substrate orientation is required[6,7].

To assess the inherent conformational propensity of different sequons we surveyed the structure of all N-x-S/T motifs where x does include P, for all human proteins in the PDB with resolution ≤ 3.0 Å, which amounts to 2,815 proteins (without duplicates) and 15,568 sequons. The results (**Fig. S9),** show that some residues, e.g. in N-**F**-S, N-**V**-T, N-**F**-T, N-**C**-T, and N-**A**-S sequons (**Fig. S9** and **S11)**, show an intrinsic secondary structure propensity that is statistically closer to the reference conformation chosen as the sequon of the peptide substrate in complex with the yeast OST from PDB 8AGE, see red dot in the Ramachandran plots (**Fig. S9**).

We expanded the secondary structure analysis to include all (human and other) proteins selected from UniRef50 and by evaluating the conformational propensity around the sequon based on the corresponding structures from the AlphaFold2 (AF2) database[8] with a confidence level pLDDT> 90, resulting in 1,822,626 sequons (**Fig. S10**). A comparison of the results between the analysis from the PDB data and AF2 data is shown (**Fig. S11**). Results are discussed below.

1. **Ramachandran plots of N-x-S/T from all human proteins in the PDB**

The analysis of the 15,568 sequons from high resolution protein structures in the PDB indicated a clear structural propensities determined by the identity of the residue at +1 and by the choice of T or S at +2. Of the 40 possible types of sequons (N-x-S = 20 and N-x-T = 20), 70% of the sequons among the 20 closest matches to the reference point (PDB 8AGE) are N-x-**T** sequons, see **Fig. S11**. This result is in agreement with the preference of T vs. S at +2 at *N*-glycosylation sites reported in previous work[9–11]. The presence of T at +2 appears to be a significant factor in determining conformational freedom of large sidechains at +1, such as W and H, but also (and much less intuitively) of A, S, M and C (**Fig. S9** and **S11**). While the structural context of all of these sequons needs to be investigated separately to understand their conformational preference, the results show clearly a structural dependence of the sequon on the structure and thus on its structural complementarity at the OST catalytic site.

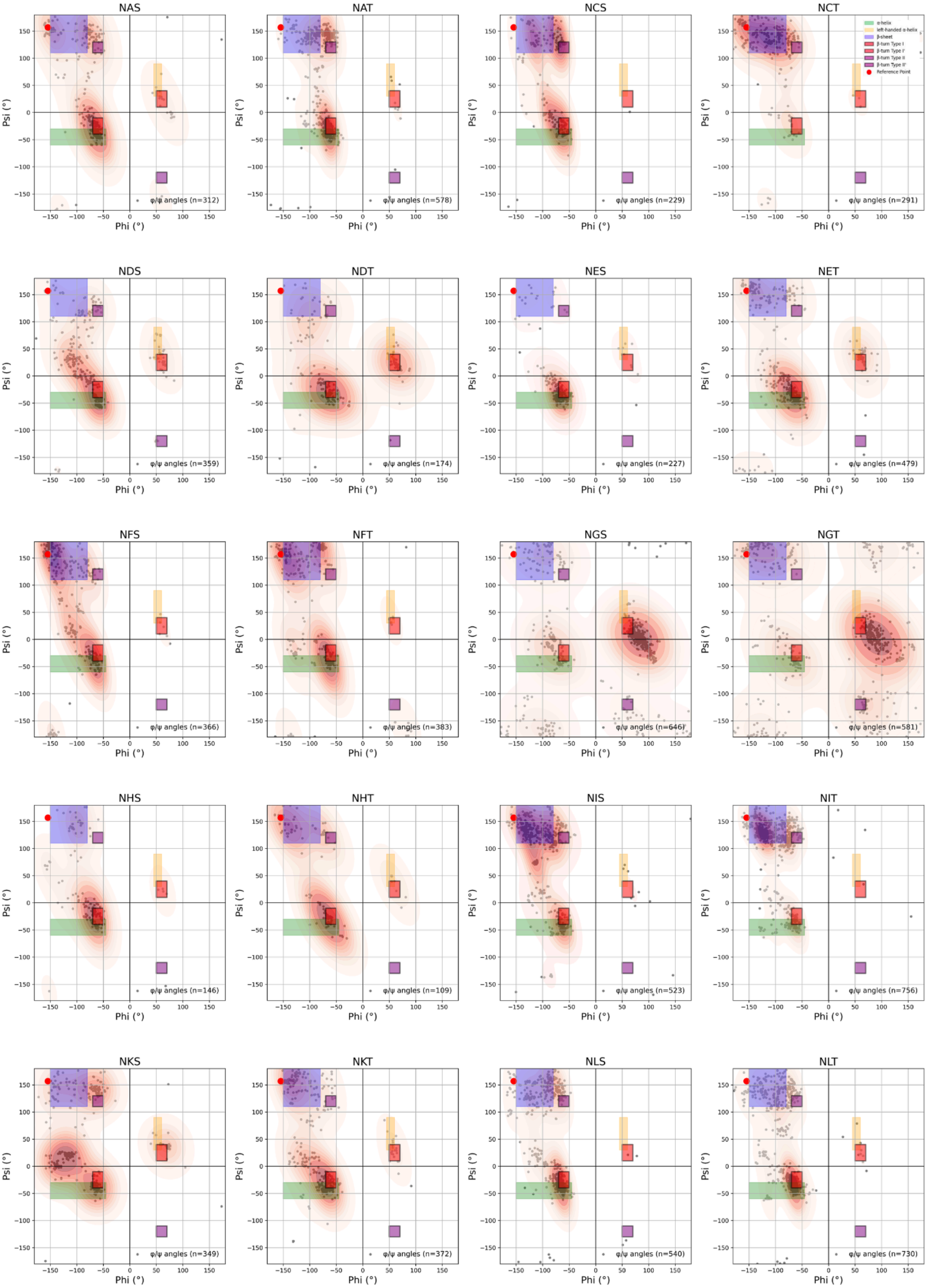
NAS
φ/ψ angles (n=312)
NAT
φ/ψ angles (n=578)
NCS
φ/ψ angles (n=229)
NCT
φ/ψ angles (n=291)
α-helix
left-handed α-helix
β-sheet
β-turn Type I
β-turn Type I'
β-turn Type II
β-turn Type II'
Reference Point
NDS
φ/ψ angles (n=359)
NDT
φ/ψ angles (n=174)
NES
φ/ψ angles (n=227)
NET
φ/ψ angles (n=479)
NFS
φ/ψ angles (n=366)
NFT
φ/ψ angles (n=383)
NGS
φ/ψ angles (n=646)
NGT
φ/ψ angles (n=581)
NHS
φ/ψ angles (n=146)
NHT
φ/ψ angles (n=109)
NIS
φ/ψ angles (n=523)
NIT
φ/ψ angles (n=756)
NKS
φ/ψ angles (n=349)
NKT
φ/ψ angles (n=372)
NLS
φ/ψ angles (n=540)
NLT
φ/ψ angles (n=730)
Phi (°)
Psi (°)

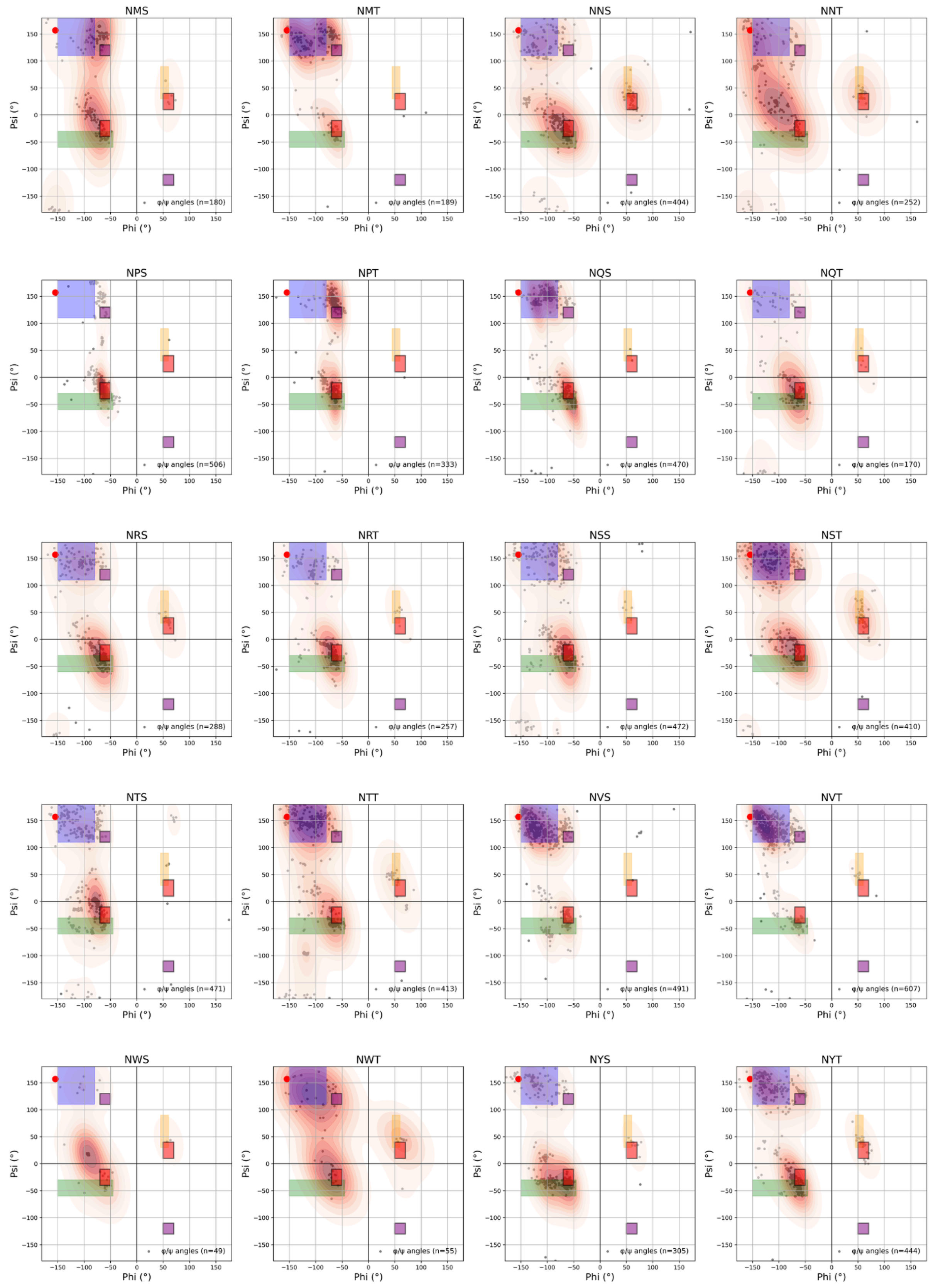


**Figure S9. Ramachandran plots of φ/ψ angles at position +1 for experimentally resolved human protein structures in the PDB with resolution ≤ 3.0 Å.** Each heat map shows the distribution of dihedral angles (gray dots) and kernel density estimates (red contours). Colored boxes indicate regions associated with secondary structures: α-helix (green), β-sheet (blue), left-handed α-helix (yellow), and β-turn types I, II, I', II' (purple). The red dot marks the φ/ψ region of the glycosylated acceptor peptide bound to yeast OST (PDB: 8AGE), used as a reference for catalytically competent conformations.

### 2. Ramachandran plots of N-x-S/T from all proteins in the UniRef50-AF2DB

The conformational propensities at the sequon obtained from the analysis of the all proteins from UniRef50 (https://www.uniprot.org/help/uniref) carrying a canonical sequon with corresponding structures from the AF2 database (AF2DB)[8] are less distinctive compared to the results from the PDB structures. The much larger number of sequons, i.e. 1,822,626 from the AF2DB vs. 15,569 from the PDB, is most certainly a determinant, which prevents a direct comparison, yet it appears that most residues are predicted to access a large part of the conformational space more readily, with the exception of Pro. While the implications of these results are interesting, finding a rationale to interpret these results is beyond the scope of this work.

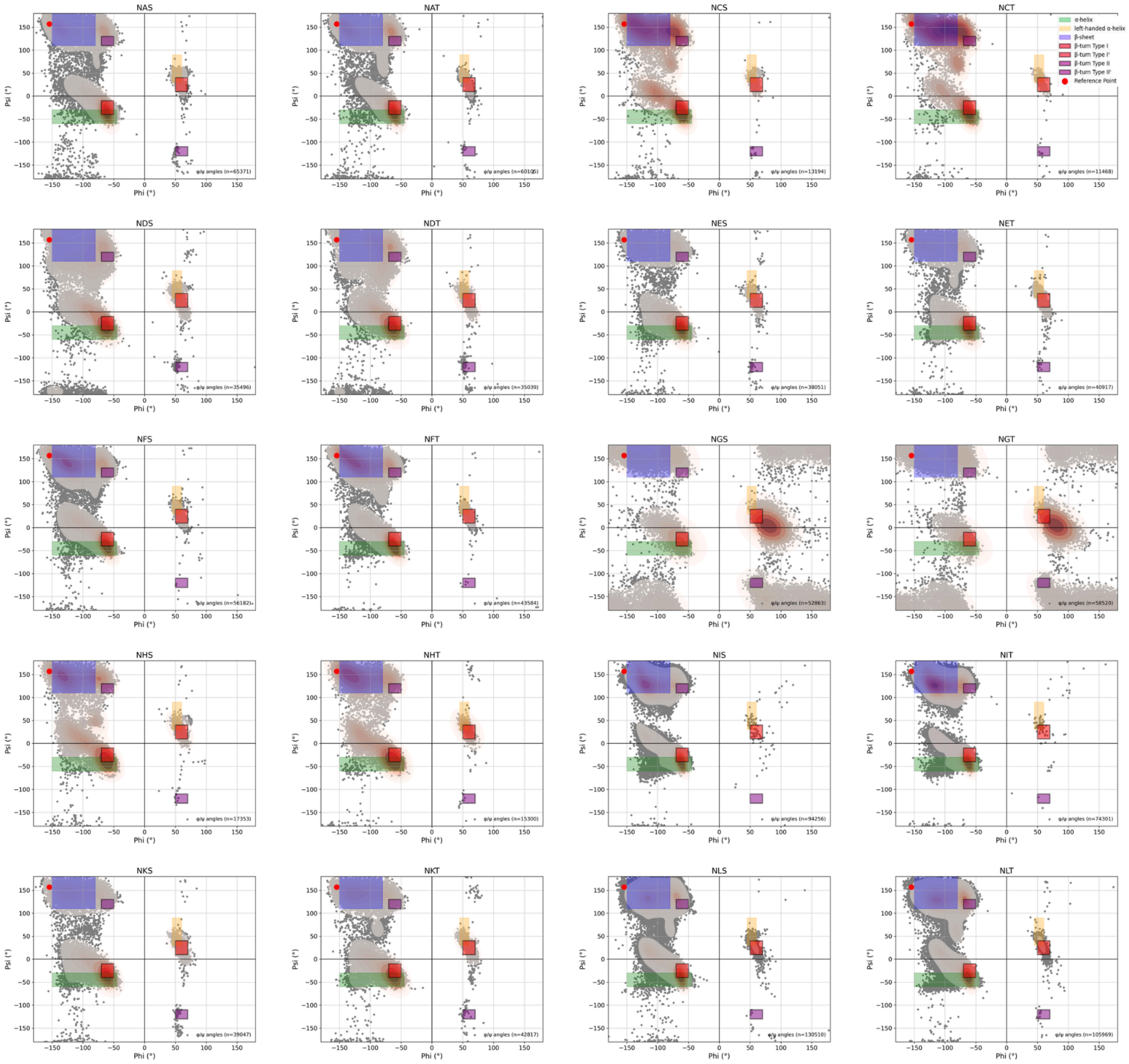

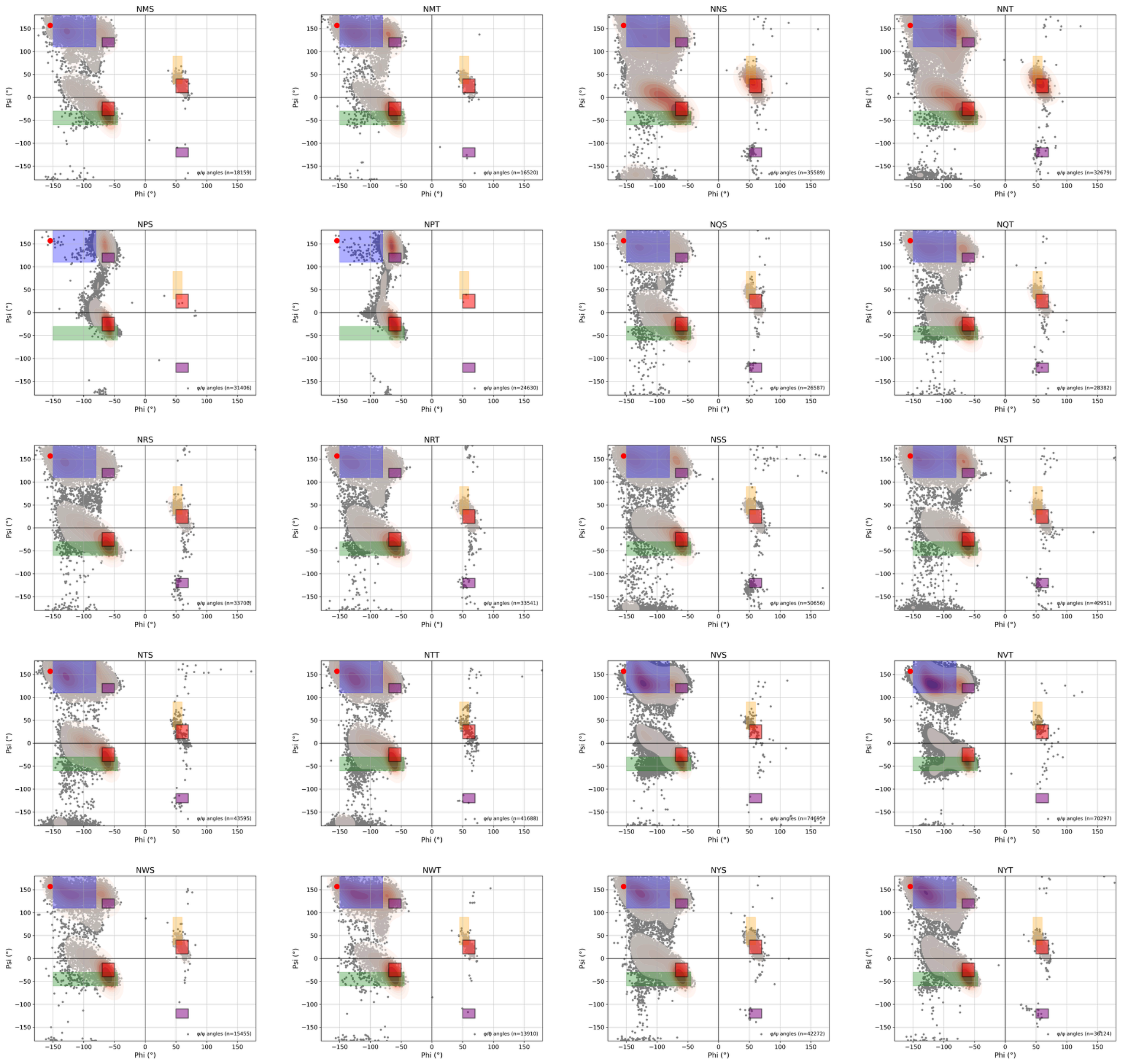


**Figure S10.** Ramachandran plots of φ/ψ torsion angles values at position +1 corresponding to AlphaFold2-predicted structures with a per-residue confidence score (pLDDT) ≥ 90 . Each heat map shows the distribution of dihedral angles (gray dots) and kernel density estimates (red contours). Colored boxes indicate regions associated with secondary structures: α-helix (green), β-sheet (blue), left-handed α-helix (yellow), and β-turn types I, II, I', II' (purple). The red dot marks the φ/ψ region of the glycosylated acceptor peptide bound to yeast OST (PDB: 8AGE), used as a reference for catalytically competent conformations.

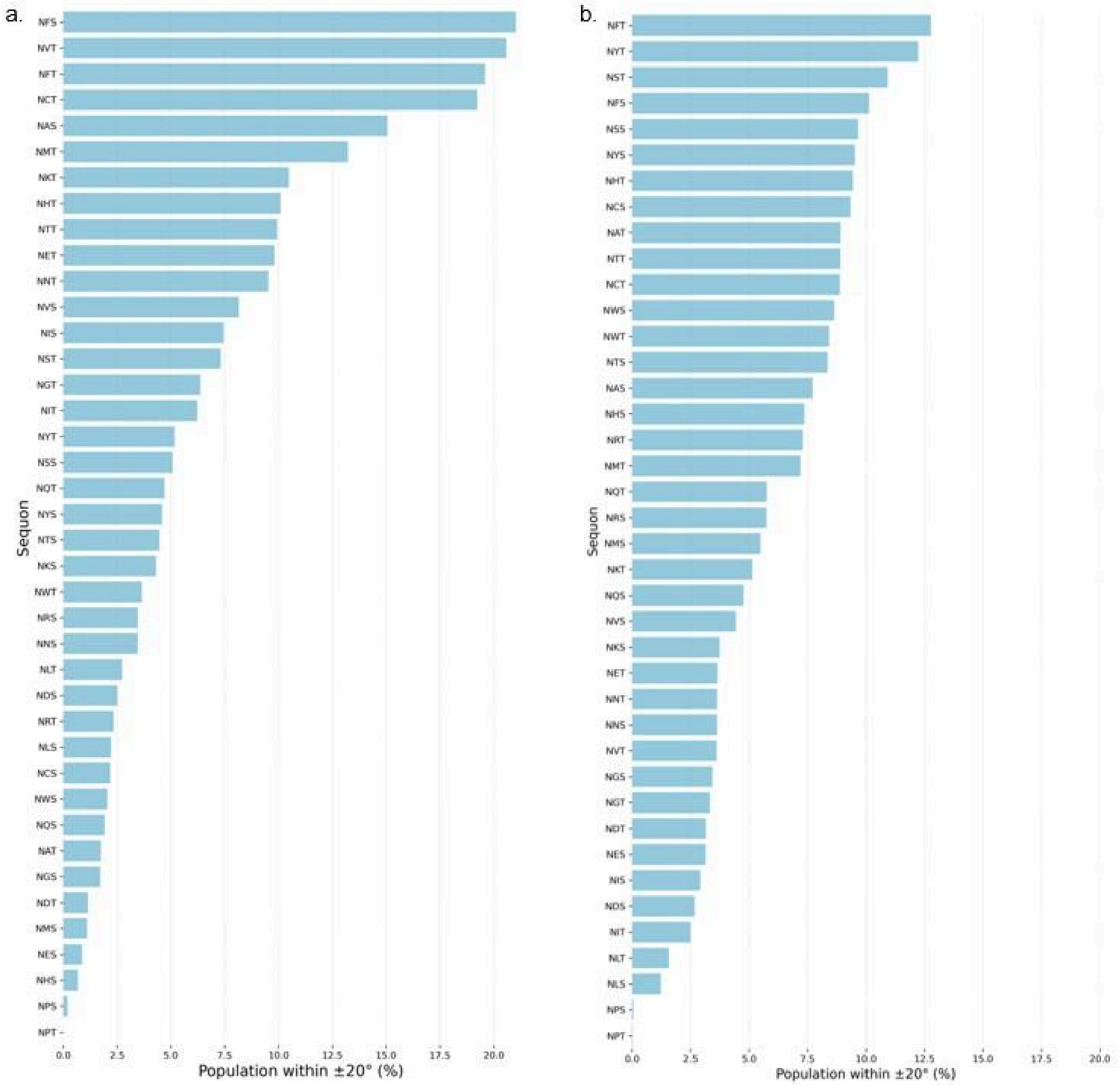


**Figure S11. Relative proportion of sequons with φ/ψ conformation within +/-20° of the reference structure in all secreted proteins (a)** in the PDB and **(b)** in the AlphaFold database.

For completeness we analysed the φ/ψ torsion angles of the peptides extracted from previously described unbiased MD simulations. Specifically, we analyzed the conformational state at the initial bound structure (at $t_0=0$ in the production trajectory) and at t = 1 μs, results are shown in heat maps (**Fig. S12**). In all simulations the φ/ψ dihedral angle values at position +1 remained within the same conformational space as the reference peptide (PDB 8AGE), both at the initial bound state and at 1 μs.

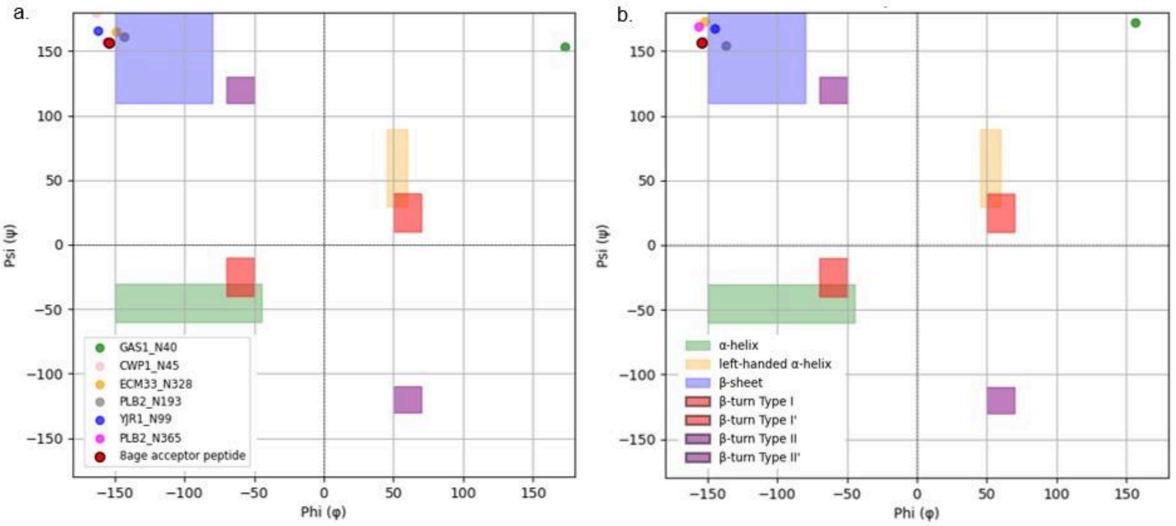


**Figure S12. Ramachandran plots of φ/ψ torsion angles at position +1 of the peptide substrates bound to the OST catalytic site. (a)** Initial bound conformations ($t_0$=0). **(b)** At t=1 μs of unbiased MD simulation. Colored dots indicate the φ/ψ values for each system (GAS1_N40, CWP1_N45, ECM33_N328, PLB2_N193, YJR1_N99, PLB2_N365, and the PDB 8AGE peptide used as reference). Colored boxes represent allowed regions for canonical secondary structures (α-helix, β-sheet, left-handed α-helix, and β-turns I/II/II′). In all cases, φ/ψ values remained confined to the same narrow region observed in the reference structure (8AGE), confirming that the extended backbone conformation at position +1 is preserved over time and is essential for stable occupancy of the catalytic pocket.

### Methods

**Yeast strains and growth conditions.** The *S. cerevisiae* yeast strains used in this study were: wild-type BY4741 (*MATa his3Δ1 leu2Δ0 met15Δ0 ura3Δ0*), *Δalg6* from the Yeast Knockout (YKO) Collection (Open Biosystems); wild-type R1158 (*MATa URA3::CMV-tTA his3Δ1 leu2Δ0 met15Δ0*) and TetO7-ALG7 (*pALG7::kanR-tet07-TATA URA3::CMV-tTA MATa his3-1 leu2-0 met15-0*), from the Tet Hughes Collection (yTHC)[1]; TetO7-ALG6 (*pALG6::kanR-tet07-TATA URA3::CMV-tTA MATa his3-1 leu2-0 met15-0*), created by insertion of the Tet-R promoter in front of the *ALG6* gene via PCR amplification and homologous recombination; TetO7-*STT3* (*MATa pSTT3::kanR-TetO7-TATACYC1 URA3::CMV-tTA his3Δ1 leu2Δ0 met15Δ0*), transformed with pRS413 encoding native or variant yeast *STT3*. DNA encoding *CWP1* was PCR amplified from yeast genomic DNA and cloned into the pRS426-GPD vector with a C-terminal His-tag. Standard techniques were used for yeast transformation and plasmid maintenance. Yeast were grown in minimal media appropriate for auxotrophic selection and with addition of antibiotics as described.

**Yeast cell wall protein preparation and mass spectrometry.** Yeast strains were grown in appropriate media to mid-log phase ($OD_{600nm}$=1) and harvested by centrifugation. Cell wall proteins covalently linked to the polysaccharide cell wall were prepared as previously described[4,12]. Briefly, cell wall-bound proteins were washed, denatured, reduced/alkylated, deglycosylated with EndoH to release *N*-glycans leaving a single *N*-acetylglucosamine (GlcNAc) at previously glycosylated Asn, and digested with trypsin. Peptides and glycopeptides were desalted with C18 ZipTips (Millipore) before LC-MS/MS analysis.

Digested peptides from yeast cell wall proteins (~1 μg) were analysed on a Waters M-Class UPLC system (running at 5 μL/min microflow) using reversed-phase chromatography. Peptides were separated on a Waters HSS T3 column (150 mm x 300 μm, 1.8 μm particles) with a 12-minute LC-MS program, with column temperature 40 ˚C. Samples were loaded in 3% pump B for 2 min, followed by a gradient of 3-45% B over 8 min, where pump A = 0.1% formic acid (FA) in water, and pump B = 0.1 % FA in acetonitrile. Eluted peptides were directly analysed on a ZenoTof 7600 instrument (ABSciex) using an OptiFlow Micro/MicroCal source. Curtain gas = 35 psi, CAD gas = 7, Gas 1 = 20 psi, Gas 2 = 15 psi, source temp = 150 ˚C, spray voltage = 5000 V, DP = 80, CE = 10. For SWATH acquisitions, an MS TOF scan across 400-1500 *m/z* was performed (0.1 sec). For MS2, variable windows spanning 398.5–1200 *m/z* were chosen for fragmentation (0.013 sec) with fragment data acquired across 140–1750 *m/z* with Zeno pulsing on. Dynamic collision energy was used.

SWATH data files were converted into DIA-NN format using DIA-NN (Version 1.8)[13]. The ion library of proteins identified was generated in DIA-NN by searching against the yeast database from UniProtKB (downloaded 20 Dec 2018) with the settings: smart profiling, out measured -rt, cleavage- K*, R*, !*P, fasta search, reannotate, reanalyse, fixed modification PPa, variable modification with HNc/ Hex/ Dea, peak centre, use quant, missed cleavage 1, threads 16, regular swath, pg label 1, original mods. The ion library generated from DIA-NN was then reformatted suitable for use in PeakView. The ion library and SWATH MS data files were imported into PeakView. Peak areas and FDR values were extracted and exported from PeakView software and analyzed as previously described[3,4,14,15].

**Computational Methods.** In our study we used the crystal structure of the ternary complex involving yeast oligosaccharyltransferase (OST), the lipid-linked oligosaccharide, and a non-acceptor peptide, TAMRA-DAB-NH2 (PDB code: 8AGC) (Ramírez et al. 2022) as starting point. To refine the protein structure, we used SwissModel (Swiss-Model Workspace et al., n.d.), incorporating the missing residues. The Ost3 lumenal domain[16] (aa 1 to 210), shown with a 3D structure obtained from AlphaFold 2[17] (Uniprot: P48439; see **Fig. 2a** in the main manuscript), was omitted from our 3D models as it is located beyond the peptide region of interest (-5 aa to +5 aa from the target Asn). Based on its location in the OST complex and orientation relative to the binding region, the Ost3 lumenal domain could be involved in tethering the substrate[18], but does not likely affect direct binding of the acceptor sequence. All complexes have a bound dolichol-diphospate linked to a $Man_3$-$GlcNAc_2$- (GlyTouCan ID G00026MO), as in the PDB structure[6]. The choice of a truncated sugar as part of the LLO, rather than the mature $Glc_3$-$Man_9$-$GlcNAc_2$-, is due to the uncertainty around the interactions of the terminal Glc residues with the OST and the consequent orientation of some of the Man residues in the arms. However, these aspects do not directly affect the acceptor's binding affinity or specificity.

All peptides were rebuilt in the OST binding site by progressively adding residues according to the sequence to the bacterial peptide in the PDB entry 5OGL (Napiórkowska et al. 2017). This peptide represents an ideal template for the peptides in our complex as its coordinates extend between +3 and -3 aa around the target Asn and it has a backbone RMSD value of 0.24 Å when aligned to the shorter substrate in the yeast OST complex (PDB 8AGC). To extract the peptide from the bacterial oligosaccharyltransferase PglB and align it correctly within our OST 3D model, we used the *super* tool in pymol to align the backbone atoms of the protein residues within 10 Å of the peptide. After structural alignment, the root mean square deviation (RMSD) value, calculated with VMD[5], between backbone atoms of the OST vs PglB atoms was 1.4 Å and the RMSD value for the peptide substrate (not included in the structural alignment) was 0.9 Å.

To reconstruct all the 31 peptides that were tested in vitro, from residue in position -4 to residue in position +4, we used ”mutagenesis” and the ”protein builder” functions available in PyMOL. From the analysis of these preliminary 3D models, we selected six complexes to explore further with MD simulations. The set covered peptides carrying two sites that are efficiently glycosylated (YJR1_N99 and PLB2_N193), two sites that are underglycosylated under stress conditions (GAS1_N40 and ECM33_N238) and two sites that are always underglycosylated (PLB2_N365 and CWP1_N45).

**Molecular Dynamic Simulations.** The 3D structure of each OST-peptide complex includes the Stt3 (residues 6–682) and Ost3 (residues 211–345) subunits, the peptide substrate (residues –10 to +10 relative to the target Asn), a dolichyl-diphosphate-linked $Man_3$ glycan donor (GlytouCan ID: G00026MO), and a $Mn^{2+}$ ion in the catalytic site.

All-atom molecular dynamics simulations were performed using version 22 of the AMBER software suite[19] with the CHARMM36m force field[20–22] Each system was embedded in a symmetric 130 Å × 130 Å lipid bilayer composed of a 1:1 mixture of POPC lipids (16:0–18:1) using CHARMM-GUI’s input generator[23,24] and solvated with TIP3P water[25] (Jorgensen et al. 1983). $Na^+$ and $Cl^-$ ions were added to neutralize the system and reach a final salt concentration of 150 mM. Protein termini were capped (ACE, CT3) and the LLO was modeled using DL2P parameters directly available in CHARMM format.

Energy minimization was performed for 5000 steps (2500 steepest descent followed by 2500 conjugate gradient), applying 10 kcal/mol·Å² positional restraints to the protein, LLO, $Mn^{2+}$ ion, peptide substrate, and lipid heads. Long range dispersion interactions were truncated with a 9 Å cutoff. The systems were gradually heated from 0 K to 310.15 K in three consecutive NVT phases of 125 ps each, with the temperature increased in stages from 0–100 K, 100–200 K, and 200–310.15 K, respectively. Temperature was controlled by Langevin dynamics with collision frequency of 1.0 $ps^{-1}$. The heating phase was followed by a 125 ps NVT equilibration at constant temperature and a 125 ps NPT equilibration at 1 bar using the Berendsen barostat[26] with semi-isotropic pressure coupling.

Three further 500 ps NPT equilibration stages were carried out, during which positional restraints were gradually reduced. In the final equilibration step, restraints of 5 kcal/mol·Å² were applied to the protein backbone, the $Mn^{2+}$ ion, and the Cα atom of the targeted Asn, while restraints of 1 kcal/mol·Å² were retained on the peptide substrate and the LLO. Total equilibration time was 2.125 ns.

All simulations were run within periodic boundary conditions where long-range electrostatics were treated with Particle Mesh Ewald (PME)[27] with a 9 Å cutoff. The SHAKE algorithm[28] was used to constrain all bonds to hydrogen atoms, enabling a 2 fs time step. Production MD simulations were performed for 1 µs per system under NPT conditions at 310.15 K and 1 bar. Weak harmonic restraints (5 kcal/mol·Å²) were maintained to restrain the glycosylation-site Asn Cα of the peptide in place. The position of the $Mn^{2+}$ ion, of the protein backbone, and of the the LLO heavy atoms were also restrained to preserve the integrity of the catalytic architecture within a reduced OST model, while enabling the sampling of the peptide conformation.

**Root-mean-square-deviation (RMSD) values.** The RMSD values of the protein Cα atoms were calculated with the *measure rmsd* command implemented in VMD with the initial coordinates of Cα atoms set as reference. The RMSD values were calculated both for the entire peptide and specifically for the sequon region corresponding to the extended binding site (-4/+4 relative to the target Asn).

**Root-mean-square-fluctuations (RMSF) values**. The RMSF values were calculated using an in-house python script with the *mdtraj* as input. The trajectories were aligned onto the initial coordinates using the Cα atoms of the entire peptide as a reference.The RMSF values were obtained for each residue along the peptides sequence.

**Potential of Mean Force (PMF) Calculations.** The PMF was calculated *via* umbrella sampling (US) simulations using the AMBER22 software package using the χ4 dihedral torsion of Arg159, defined by atoms CG, CD, NE, and CZ, as reaction coordinate. The initial structure for umbrella sampling was selected from a well-equilibrated frame of the same system used in the unbiased simulations. Positional restraints applied during umbrella sampling were kept consistent with those used in the unbiased simulations. Sampling covered the 0°–360° range using 181 windows spaced every 2°. A harmonic restraint of 200 kcal/mol·rad² was applied to χ4 in each window. Each window included energy minimization (5000 steps), NPT equilibration (500 ps at 310 K and 1 atm, Langevin thermostat, SHAKE, semi-isotropic pressure coupling), and 5 ns production MD with χ4 recorded every 2 ps. The total sampling was 995.5 ns for each system.

We reconstructed the PMFs using the Weighted Histogram Analysis Method (WHAM)[29], which combines biased distributions and removes the restraint contribution to recover the unbiased free energy landscape. Since AMBER defines the restraint as $U(x) = k(x - x_0)^2$ in kcal/mol·rad², and WHAM uses $U(x) = ½k(x - x_0)^2$ in kcal/mol·deg², we converted the force constant by applying the factor $2 \times (\pi/180)^2$, yielding 0.12184 kcal/mol·deg², applied uniformly across all windows. Convergence was evaluated using WHAM by examining PMF stability across iterations and histogram overlap between adjacent windows.

**Secondary structure analysis and data mining.** To determine the sequence composition of internal (not solvent accessible) Asn within sequons, we first selected from the initial set of 3,648 reviewed secreted human proteins retrieved from UniProtKB all 14,835 canonical NXS/T sequons, by filtering the search for proteins with taxonomy ID 9606 (*Homo sapiens*) and subcellular location SL-0243, which corresponds to secreted proteins. Of these only 997 sequons (8%) had a relative solvent accessibility (RSA) value at the Asn $< 0.15$ and were classified as buried, corresponding to 567 unique proteins. Residue enrichment was monitored across the –4 to +4 window around the Asn target by running a global pairwise alignment using the *pairwise2.globalxx* function in Biopython[30]. We calculated enrichment values for each position in the –4 to +4 region by dividing the observed frequency of each amino acid by its expected background frequency in the vertebrate proteome[31]. We identified the following residues enriched within the -4/+4 window IGEI**N**H[S/T][W/G]N. Notably this sequence should be interpreted as a list, as each position is independently counted.

To analyze the conformational (secondary structure) preferences around canonical sequons, we computed φ (phi) and ψ (psi) dihedral angles for all human canonical sequons for which structural data were available in the Protein Data Bank (PDB) with a resolution ≤ 3 Å for a total of 15,569 sequons. We augmented this structural analysis by considering sequons in all proteins in the UniRef50 dataset with structures from AF2DB corresponding to a per-residue confidence level pLDDT ≥ 90 across the –4 to +4 window (1,822,626 sequons).

Analogously, one could correctly consider non-canonical sequons as an example of optimal substrates for *N*-glycosylation. We were able to collect only 167 experimentally-evidenced examples of *N*-glycosylated non-canonical sequons. This data set is unfortunately too limited for the analysis to be conclusive, with the only significant feature being the presence of Cys in almost all sequences, and at all positions. More work in this area is currently ongoing.

**Recombinant Fab production and purification methods.** Fabs containing PNGS for occupancy analyses were co-transfected into MEXi HEK-293E cells (IBA-LifeSciences) at a 1:1 molar ratio, following the IBA LifeSciences protocol. Transfections were performed on a 30 ml to 200 ml scale using a DNA:PEI Max ratio of 1:3. After incubating the cells at 37 °C for seven days, the supernatants were harvested and passed through a 250 ml Stericup-HV 0.22 µm sterile filter (Millipore). Lymphoma-derived F(ab)s were then purified from the supernatants using a 5 ml CaptSelect CH1-XL affinity matrix column on an NGC medium-pressure liquid chromatography system (Bio-Rad). The elution fractions were pooled and concentrated to 500 µl using Amicon® Ultra-15 centrifugal filter devices (Merck). Subsequently, the Fabs were further purified into PBS with a Superdex 200 Increase 10/300 GL gel filtration column (Cytiva). The column was pre-equilibrated in PBS, and the purified Fab material was injected into the NGC chromatography system. Fractions were collected and pooled according to their corresponding peaks on the gel filtration profile.

**Glycopeptide mass spectrometry of Fabs.** Individual 50 µg aliquots of purified F(ab)s were digested with trypsin, chymotrypsin (Promega), and/or α-lytic protease (NEB) at a 1:30 (w/w) enzyme-to-substrate ratio in 50 mM Tris/HCl, pH 8.0, to generate glycopeptides containing a single N-glycosylation site. Resulting glycopeptides were purified using C18 Zip-tips (Merck Millipore), dried, resuspended in 0.1% formic acid, and analyzed by LC-MS using an Ultimate U3000 system coupled to an Orbitrap-Eclipse mass spectrometer (Thermo Fisher Scientific). Fragmentation of peptide and glycopeptide ions was performed by stepped higher-energy collisional dissociation. Separation was carried out on an EasySpray PepMap RSLC C18 column (75 µm × 75 cm) using a 0–32% acetonitrile gradient in 0.1% formic acid over 240 minutes, followed by 80% acetonitrile in 0.1% formic acid for 35 minutes (flow rate: 200 nl/min; spray voltage: 2.5 kV; capillary temperature: 275 °C). Glycopeptide fragmentation spectra were processed with Byos v3.5 (Protein Metrics Inc.), and the relative abundance of each glycoform at individual sites was quantified.

**Crystallographic Data Collection and Analysis**

**Table S2.** Crystallographic data collection and refinement statistics for Fab 10531. Values for the highest resolution shell are reported in parentheses.

| | |
|---|---|
| Data Collection | |
| Beamline | ID30B (European Synchrotron Radiation Facility) |
| Resolution range (Å) | 49.8–1.76 (1.79–1.76) |
| Space group | $P2_12_12_1$ |
| Unit cell dimensions:<br>*a, b, c* (Å)<br>a, b, g (°) | <br>59.5, 67.7, 149<br>90, 90, 90 |
| Wavelength (Å) | 0.89 |
| Unique reflections | 60,669 (2,953) |
| Completeness (%) | 100 (100) |
| $R_{merge}$ | 0.101 (1.29) |
| $R_{meas}$ | 0.105 (1.34) |
| $R_{pim}$ | 0.029 (0.363) |
| $I$/s($I$) | 12.0 (0.60) |
| Multiplicity | 13.6 (13.5) |
| $CC_{1/2}$ | 0.999 (0.689) |
| Wilson *B* factor ($Å^2$) | 27.4 |
| Refinement | |
| Number of reflections (all / free) | 60,594 / 3,007 |
| $R_{work}$ (%) | 17.8 |
| $R_{free}$ (%) | 21.2 |
| RMSD[1]:<br>Bonds (Å)<br>Angles (°) | <br>0.0091<br>1.65 |
| Molecules per ASU[2] | 1 |
| Atoms per ASU | 3,947 |
| Average *B* factors ($Å^2$) (protein / carbohydrate / water) | 39.2 / 70.2 / 48.6 |

| | |
|---|---|
| Model quality (Ramachandran plot)[3]: | |
| Most favoured region (%) | 93.1 |
| Allowed region (%) | 5.07 |
| Outliers (%) | 0.92, 0.00[4] |

[1]RMSD, root-mean-squared-deviation

[2]ASU, asymmetric unit

[3]as calculated with Molprobity[32]

[4]as reported by the wwPDB Structure Validation Report

**Crystallisation and structure solution of Fab 10531.** Fab 10531 was exchanged into a buffer comprising 50 mM HEPES and 150 mM KCl (pH 7.5) prior to crystallization, using a Vivaspin 20 centrifugal concentrator (MWCO 10 kDa; Cytiva). Sitting drop vapour diffusion crystallisation experiments were set up with an Oryx4 robot (Douglas Instruments), which yielded crystals of Fab10531 in a condition comprising 0.09 M halogens, 0.1 M buffer system 2 (pH 7.5) and 37.5 % v/v precipitant mix 4 (condition B8 within the Morpheus crystallization screen[33]; Molecular Dimensions). Crystals were flash-frozen and stored in liquid nitrogen prior to data collection (additional cryoprotection was not required, due to the presence of cryoprotectant inherently within the mother liquor).

Data collection was carried out on beamline ID30B at the European Synchrotron Radiation Facility (under proposal mx2639). Diffraction data was collected from crystals placed within a 100 K cryostream and using an X-ray wavelength (λ) of 0.886 Å. Diffraction images were processed using DIALS[34]. The structure of Fab 10531 was solved to 1.76 Å by molecular replacement (using the program Molrep[35]) within the ccp4i2 software suite[36]), using an AlphaFold 3[37]) model of the Fab as the search model. This initial solution was improved with successive rounds of model building and refinement, using programs Coot[38,39] and Refmac5 [8], respectively. Structure refinement was performed using automatically-generated translation/libration/screw restraints in Refmac5[40]. Glycan conformations were validated using Privateer[41]). The model was validated using MolProbity[32] and the PDB validation server[42] prior to deposition. Statistics for data collection and refinement of the Fab 10531 structure are presented within **Table S2**.

PDBePISA[43] was used to calculate the solvent accessibility of residue N57 in the Fab 10531 structure. Figures depicting the protein structure were generated using UCSF ChimeraX[44].
